# Myofibroblast differentiation is governed by adhesion mechanics, and inhibition of Talin2 reverses lung and kidney fibrosis

**DOI:** 10.1101/2021.06.07.447403

**Authors:** Michael JV White, Melis Ozkan, Jorge Emiliano Gomez Medellin, Ani Solanki, Jeffrey A Hubbell

## Abstract

Fibrosis is involved in 45% of deaths in the United States, and no treatment exists to reverse progression of the disease. In order to find novel targets for fibrosis therapeutics, we developed a model for the differentiation of monocytes to myofibroblasts that allowed us to screen for proteins involved in myofibroblast differentiation. Inhibition of a novel protein target generated by our model, talin2, reduces myofibroblast morphology, α-smooth muscle actin content, collagen I content, and lowers the pro-fibrotic secretome of myofibroblasts. We find that knockdown of talin2 de-differentiates myofibroblasts, talin2 knockdown reverses bleomycin-induced lung fibrosis in mice, and Tln2 -/-mice are resistant to unilateral ureteral obstruction-induced kidney fibrosis and are resistant to bleomycin-induced lung fibrosis. Talin2 inhibition is a potential treatment for reversing lung and kidney fibroses.

**One Sentence Summary:** Silencing the stress sensor Talin2 reverses myofibroblast differentiation and existing fibrosis.

## INTRODUCTION

Fibrosing diseases—including pulmonary fibrosis, congestive heart failure, liver cirrhosis, and end-stage kidney disease—are involved in 45% of deaths in the United States [1, 2]. There are few FDA approved treatments for fibrosis [2, 3]. Currently approved FDA treatments (pirfenidone and nintedanib) slow, but do not reverse, the progression of fibrosis [4]. Further, the mechanisms of action of pirfenidone and nintedanib are poorly understood [5]. To date, only one treatment (recombinant pentraxin-2, PRM-151) has shown even a modest ability to reverse fibrosis in some patients [6].

The ultimate goal of any treatment is to reverse fibrosis [1]. Interrupting collagen deposition destabilizes scar tissue and is a necessary prerequisite for reversing fibrosis [1]. Another prerequisite for reversing fibrosis is removing deposited ECM while regenerating tissue, of which monocyte-derived cells are capable [7].

Myofibroblasts are key to fibrosis progression, but the term “myofibroblast” denotes the function of a cell rather than a precise cell type [8]. Thus, myofibroblasts can arise from several progenitors, including hepatocytes, fibroblasts, epithelial cells, and monocytes [1, 8, 9].

Myofibroblasts promote scar tissue by actively depositing ECM, secreting pro-fibrotic signals, and by physically stiffening tissues by tension force generated by their cytoskeleton [1, 10] [11]. Myofibroblast contractility is key to the progression of fibrosing diseases, and tissue stiffness can activate myofibroblast phenotypes independent of TGF-β [12]. Myofibroblasts are formed or recruited in response to injury [9]. Normal, undamaged tissues “stress-shield” cells from tension, with intact ECM supporting the stress. By reducing the stiffness of the environment, normal healthy tissues are free of myofibroblast activation [13]. Among myofibroblast precursors, monocytes are unique in that they can be recruited from the bloodstream to different tissues in the body [14]. These monocyte-derived myofibroblasts may provide a third of myofibroblasts in liver fibrosis [15], kidney [16], lung [17, 18], and skin fibroses [19].

Myofibroblast presence in damaged tissue can either be resolved, in the case of scarless wound healing, or can lead to the formation of scar tissue [11]. As myofibroblasts are key to scar tissue formation and maintenance, reversing fibrosis will involve de-activating or removing myofibroblasts from scar tissue [11]. It has been suggested that apoptosis [20] or de-differentiation [9] are the two main methods that the body de-activates myofibroblasts [21].

Mechanosensing is key to maintenance of the myofibroblast phenotype [11]. Mechanosensing is a dynamic process integrating multiple signals between cell surface integrins, intracellular machinery, and secreted signals [10]. In mechanosensing, changes in interactions between integrins, intracellular proteins, and the actin cytoskeleton can lead to signaling changes in what might otherwise appear as a static environment [22]. Among these changes are the maturation of focal adhesions (FAs), which are a dynamic, responsive link at the cell surface between the actin cytoskeleton and the ECM [22, 23]. Mechanoproteins interact in FAs, leading to a complex signaling and force-distribution environment [24]. FAs exist in a dynamic state that ranges from transient—in migrating cells—to super-mature in some myofibroblasts [25].

Fibrillar adhesions (FBs) are a complex of cytoskeletal machinery and actin that connect to FAs and localize into long intracellular bundles [26, 27]. Talin is an intracellular tension-sensing adapter protein that is key to FAs and FBs [26]. Talin has two isoforms (1 and 2) that share ∼76% homology and are both 270 kDa [28]. Talin is composed of an N-terminal head domain and a C-terminal rod domain [29, 30]. Each talin has different expression patterns in different tissues [28, 31, 32]. Talin1 and talin2 have distinct but overlapping interactions [33-35], cellular localizations [26, 33], and functions [33, 35]. Talin2 is sometimes capable of rescuing phenotypes caused by talin1 knockout, and vice-versa [36].

Talin2 has splice-variants, though the function of these has not yet been determined [28, 31]. Talin1 is ubiquitously expressed in vertebrates, and the knockout is embryonic lethal [37]. Talin2 is expressed only in certain tissues, and knockout mice are either asymptomatic [38], or mildly dystrophic at advanced ages [28].

Talins are key to mechanosensing and mechanotransduction [33, 35]. Talins function as adapter proteins, classically explained as binding integrins with their head domains, and binding vinculin and other FA and FB proteins with their rod domains [30, 39-42]. Recent studies have shown that talins interact more broadly with the adhesome, as talin’s individual domains each have affinities of their own for integrins or cytoskeletal proteins [33, 36, 43, 44]. Talins provide a framework on which all components of focal adhesions are built [33].

Here we show that monocytes and fibroblasts only differentiate into myofibroblasts when adhered to a surface of sufficient stiffness. We were motivated to target talin2 by an RNAseq (mRNA sequencing) study comparing gene expression of myofibroblasts cultured on prof-fibrotic, sufficiently stiff surfaces versus culture on anti-fibrotic, insufficiently stiff surfaces.

Knockdown of talin2 interrupts the ability of monocytes to sense stiffness, prevent monocyte-myofibroblast differentiation, and de-differentiates myofibroblasts. We further show that knockdown of talin2 may have therapeutic utility in pulmonary fibrosis.

## RESULTS

### Myofibroblast differentiation is governed by adhesion and substrate stiffness

We began our studies investigating the features of cell adhesion that lead to efficient differentiation of monocytes into myofibroblasts, seeking to identify a key pathway that could be inhibited to prevent differentiation. This led to the identification of a key role of talin2, knockdown of which both prevented differentiation of monocytes into myofibroblasts and moreover dedifferentiated existing myofibroblasts. We then investigated talin2 knockdown in a murine model of pulmonary fibrosis, which confirmed a key role of talin2.

Our results demonstrate that monocyte differentiation into myofibroblasts is governed by the cell adherent state, including the elastic modulus of the adhesion substrate. First, when exposed to pro-fibrotic factors (tryptase, IL-13) for 1 hr, human monocytes in suspension and later cultured adherently do not differentiate efficiently into myofibroblasts (as determined by morphological demonstration of a clear spindle shape), compared to monocytes exposed to the same pro-fibrotic factors for the same amount of time while adhered (Fig 1A). Adhered monocytes also increase their double-positive αSMA and collagen I content (Fig 1B), when compared to suspended-then-adhered monocytes, as determined by flow cytometry.

**Figure 1:**
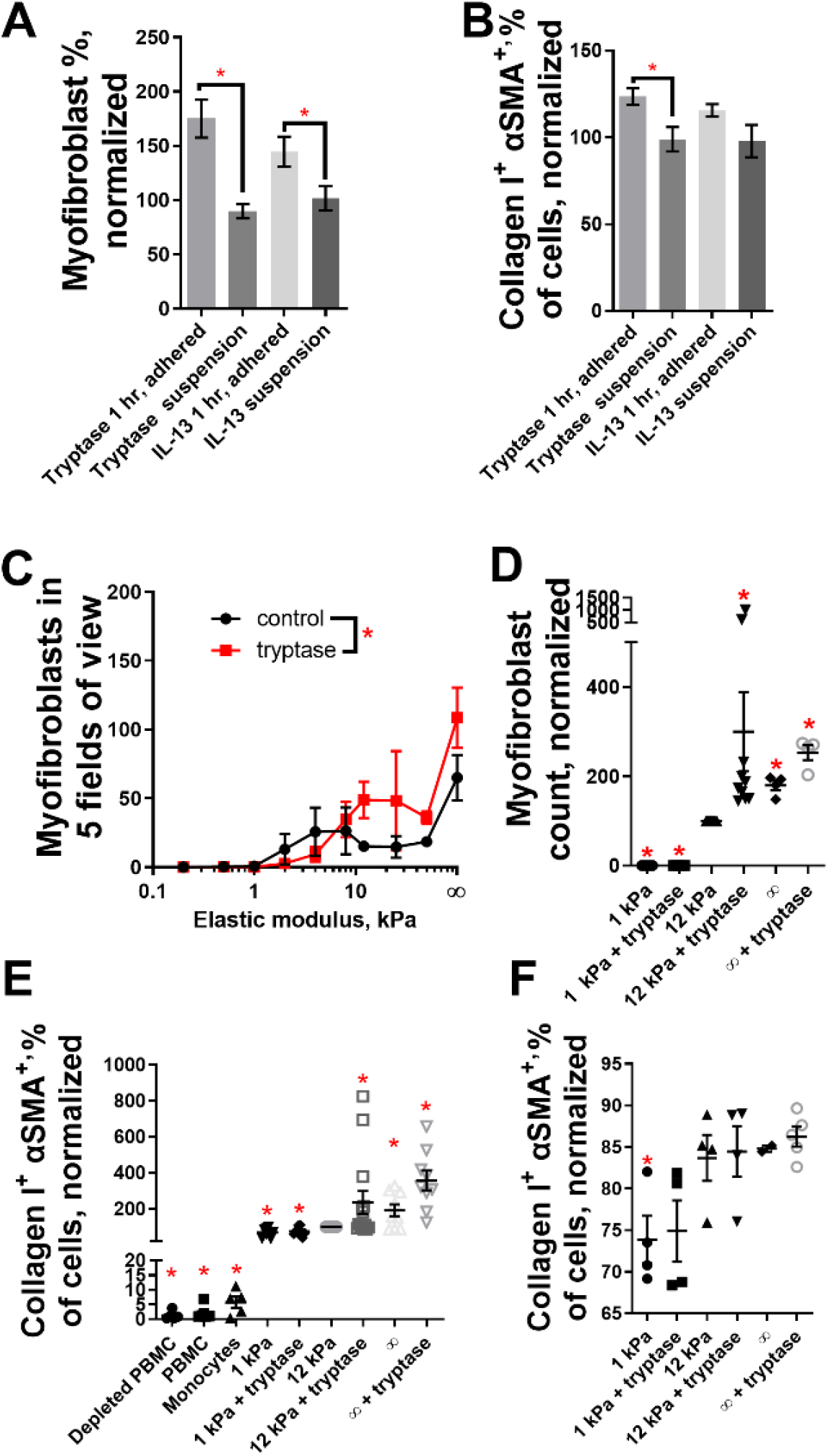
Monocyte-to-myofibroblast differentiation requires multiple adhesion-related checkpoints to proceed. (A) Freshly isolated human monocytes were cultured with or without pro-fibrotic factors (tryptase, IL-13) for 1 hr, either adhered to a tissue-culture treated surface or in suspension, and then differentiated in fresh medium lacking the pro-fibrotic factors over 3 days into myofibroblasts adherent to the tissue-culture treated surface. Differentiation into myofibroblasts was determined by morphology, namely a highly-elongated, spindle shape. Due to donor variability, readouts for all human monocytes are normalized and compared to the same individual donor’s myofibroblast numbers, αSMA and collagen I content. (B) Myofibroblasts were then removed from the surface and assessed for the percentage of αSMA-and collagen I-positive cells by flow cytometry, again normalized for each individual donor. (C) Monocytes plated on fibronectin-coated surfaces differentiated into myofibroblasts, with or without a pro-fibrotic factor, more on stiffer than on softer surfaces. Monocytes cultured at 1 kPa, 12 kPa, and on tissue culture treated plastic (functionally infinite kPa), each coated with fibronectin, form myofibroblasts at increasing amounts, as measured by (D) morphology (normalized to 12 kPa values, statistics vs 12 kPa) and (E) percentage of cells that are αSMA and collagen I positive by flow cytometry (normalized to 12 kPa values, statistics vs 12 kPa). Due to donor cell variability, readouts are normalized and compared to the number of positive cells at 12 kPa for the individual donor. Donor PBMC, isolated monocytes, and PBMC depleted of monocytes were assessed by flow cytometry and normalized to the 12 kPa readouts as well. Statistics are compared to 12 kPa. (F) Surfaces of 1, 12, and functionally infinite kPa stiffness induce human fibroblast cultures to become increasingly αSMA- and collagen I-positive by flow cytometry. n ranges from 3 to 20. * = statistical significance of P < 0.05, < 0.01, or < 0.001, Statistical comparisons are to the 12 kPa value for each dataset (D-F), 2-way ANOVA for panel C, Sidak post-test, Student’s t-test for other panels.

Second, human monocytes are unable to differentiate into myofibroblasts on surfaces softer than 1 kPa even when adherent; monocyte-myofibroblast differentiation can occur and be potentiated at 12 kPa (Fig 1C) in the presence of a pro-fibrotic factor. For reference, Supplemental Table 1 (data from [45], [46]) shows the stiffnesses of various human tissues.

Culture on surfaces that had been pre-coated with the ECM protein fibronectin supported myofibroblast differentiation even without pre-treatment with a pro-fibrotic factor, yet in a manner that is dependent on the substrate stiffness, with monocytes cultured on 1, 12, and functionally infinite kPa fibronectin-coated surfaces showing increasing amounts of myofibroblast differentiation (Fig 1D), including increasing percentages of cells double-positive for αSMA and collagen I (Fig 1E). Cultured fibroblasts also increase the percentage of αSMA-and collagen I-double-positive cells on higher surface stiffness fibronectin-coated substrates, both in the absence and presence of pro-fibrotic factors (Fig 1F).

### Talin2 is upregulated in myofibroblasts cultured on stiff surfaces

Using these results (Fig 1C-E) as a guide, we cultured human monocytes from three donors on 1 and 12 kPa fibronectin-coated surfaces, isolated total mRNA from the population, and performed an RNAseq investigation (Supplemental tables 2 and 3). This RNAseq analysis revealed genes that were differentially expressed between monocytes cultured at 1 and 12 kPa, yielding both down-regulated and up-regulated individual genes and pathways. Individual upregulated genes included collagens (e.g., collagen XXII upregulated 22-fold) and chemotactic factors (CCL22 upregulated 13-fold). However, analysis of upregulated pathway allowed for a more inclusive and complete picture of the changes induced by culturing monocytes on stiffer surfaces under pro-fibrotic conditions. Specifically, upregulated pathways related to cell adhesion included the paxillin and integrin pathways. Among the individual genes that were downregulated at 1 kPa (not supporting myofibroblast differentiation) relative to 12 kPa (supporting), we noticed the stress sensor talin2, which was reduced by 3-fold on the surface that did not allow monocyte-to-myofibroblast differentiation.

Based on this observation, we focus the remainder of this study on this intracellular modulator of adhesion mechanics, the tension-sensing protein talin2. We explore the potential of integrins to modulate myofibroblast differentiation in a companion study [47].

To confirm that the measured mRNA reduction in the sequence for talin2 at low stiffness corresponded to a drop in the protein talin2, we cultured monocytes and fibroblasts on surfaces of increasing stiffness in the presence and absence of a pro-fibrotic signal. Expanding on the staining for collagen I and αSMA in Figs 1E and 1F, talin2 was low at 1 kPa and increased in both human monocytes (Fig 2A) and human fibroblasts (Fig 2B) on stiffer surfaces in more pro-fibrotic environments.

**Figure 2:**
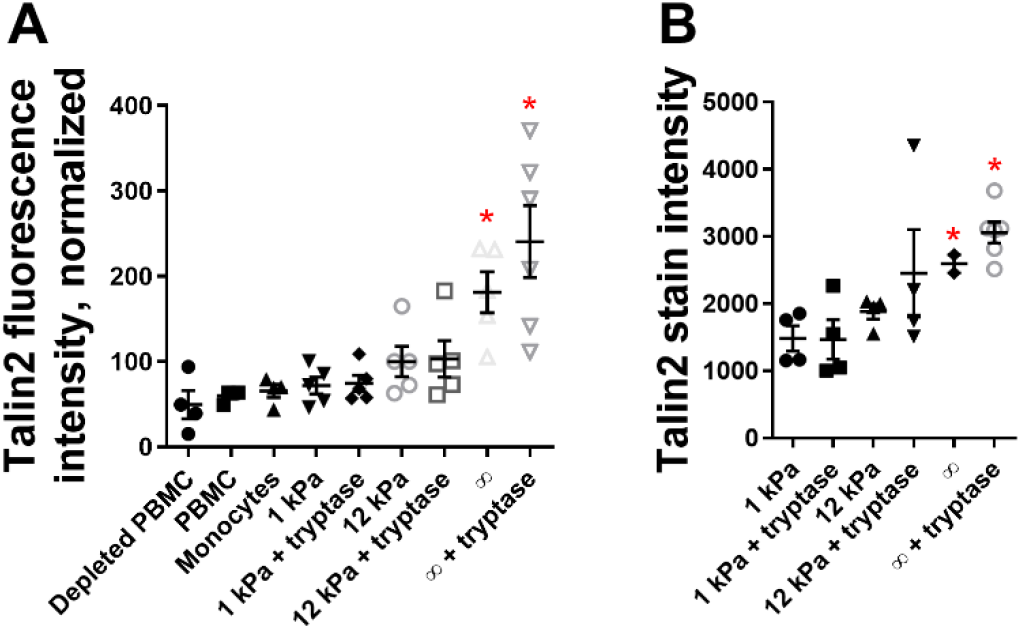
Increasing surface stiffnesses and pro-fibrotic conditions increase the intensity of talin2 immunostaining in human cells. (A) Freshly purified monocytes, and (B) cultured fibroblasts, were cultured conditions of increasing stiffness and in the presence of pro-fibrotic factors. n ranges from 2 to 8. * = statistical significance of P < 0.05, < 0.01, or < 0.001, Statistical comparisons are to the 12 kPa value for each dataset., Student’s t-test.

To determine if mouse cells also increase talin2 in more pro-fibrotic environments on stiffer surfaces, we cultured monocytes purified from mouse spleens and mouse fibroblasts under conditions similar to those used in our study of human cells reported in Figs 1 and 2. Mouse monocytes differentiated into morphologically spindle-shaped myofibroblasts with increasing surface stiffness (Fig 3A), increasing positivity for αSMA and collagen I staining (Fig 3B), and increasing talin2 staining (Fig 3C). Mouse fibroblasts also increase in αSMA and collagen I staining (Fig 3D) and talin2 staining (Fig 3E) on stiffer surfaces in presence of tryptase.

**Figure 3:**
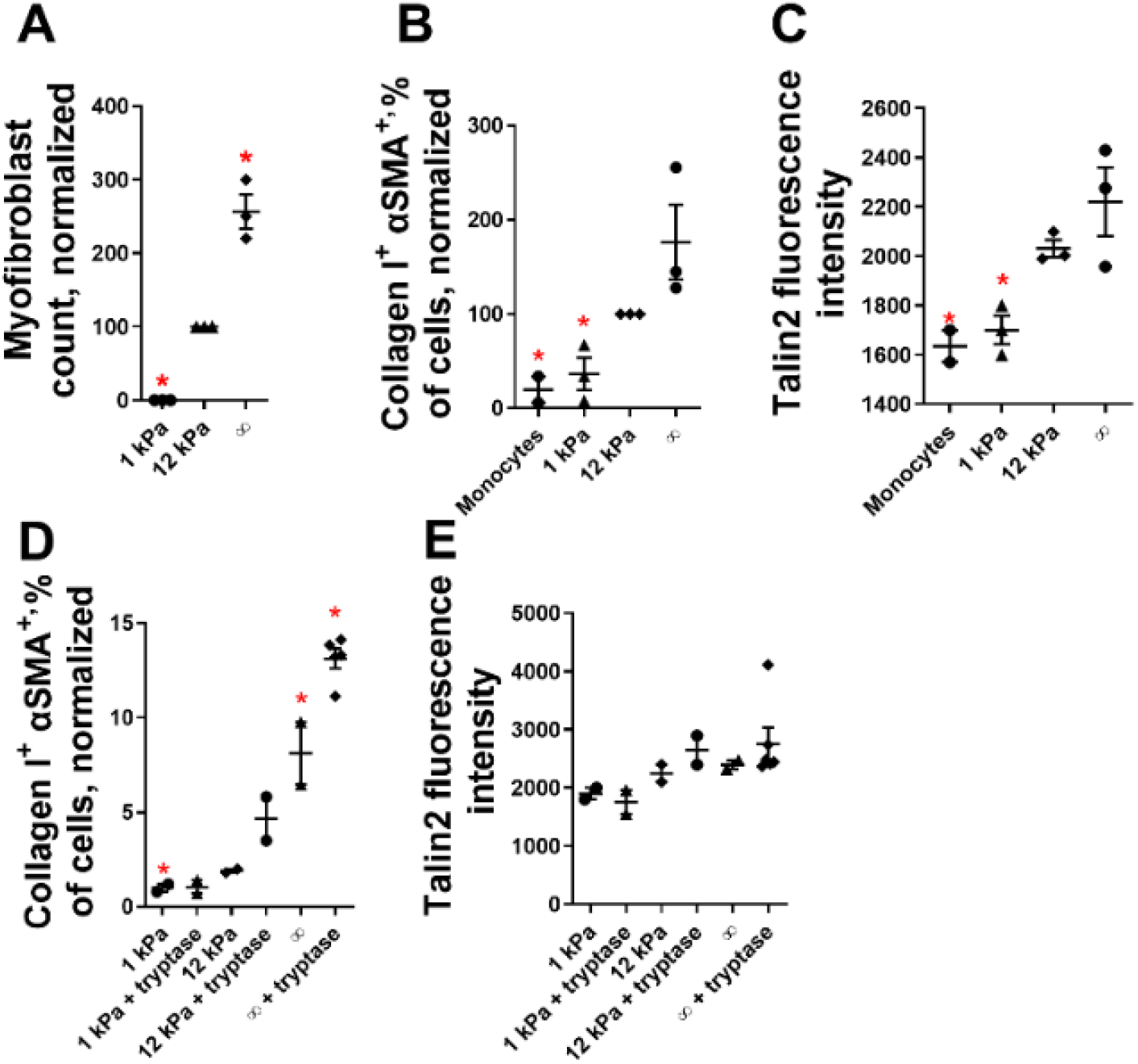
Increasing surface stiffnesses increases myofibroblast differentiation and the intensity of talin2 immunostaining in mouse cells. Freshly purified mouse monocytes cultured at 1, 12, and functionally infinite kPa form myofibroblasts at increasing amounts, as measure by (A) morphology and (B) percentage of cells that are αSMA and collagen I positive. (C) These same populations show increasing talin2 staining intensity. Mouse fibroblasts cultured under identical conditions also show increasing numbers of (D) cells that are αSMA and collagen I positive and (E) increased talin2 staining intensity. n ranges from 2 to 8. Statistical comparisons are to the 12 kPa value for each dataset. * = statistical significance of P < 0.05, < 0.01, or < 0.001, Student’s t-test.

### Inhibition of Talin2 reverses myofibroblast differentiation and existing fibrosis

To determine if inhibition of talin2 expression could de-differentiate myofibroblasts, we allowed human and mouse monocytes to become myofibroblasts, and treated those myofibroblasts with a mixture of 4 non-targeting silencing RNAs (siRNA), 4 human talin2 siRNAs, and 4 mouse talin2 siRNAs. The mouse and human talin2 siRNA mixtures share one sequence in overlap. To establish a dose range, we treated human monocyte-derived myofibroblasts with the talin2 siRNA mixture, yielding an IC50 of 15 nM (Fig 4A) for inhibition of myofibroblast morphology. Treatment with 50 nM of talin2 siRNA also reduced the percentage of αSMA and collagen I positive human monocytes (Fig 4B), more than the control siRNA mixture. Only the human talin2 siRNA significantly reduced the amount of talin2 (Fig 4C) for human monocyte-derived myofibroblasts.

**Figure 4:**
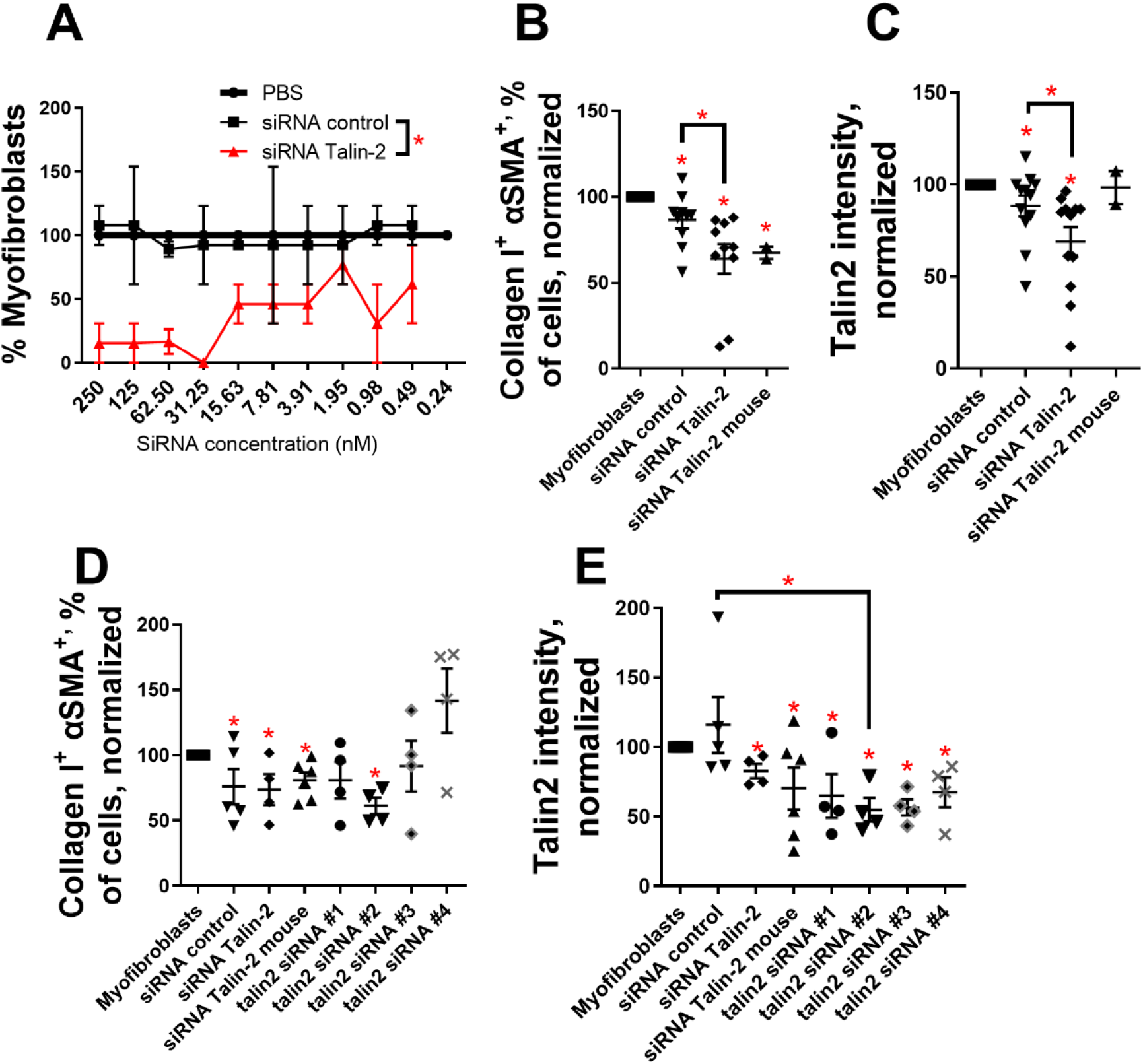
Silencing RNA against talin2 dedifferentiates human and mouse myofibroblasts. Freshly purified human monocytes allowed to become myofibroblasts were treated with (A) a mixture of 4 non-targeted siRNAs and talin2 siRNAs at the indicated concentrations. (B) myofibroblasts differentiated from human monocytes were treated with 50 nM mixtures of control siRNA, and siRNA targeting human or mouse talin2, and assessed for (B) the number of αSMA^+^ Collagen I^+^ cells and (C) the amount of talin2. Myofibroblasts differentiated from mouse monocytes were treated identically, except that individual siRNA was also used in addition to the mixtures, and were assessed for (D) the number of αSMA^+^ Collagen I^+^ cells and (E) the amount of talin2. n ranges from 2 to 10. * = statistical significance of P < 0.05, < 0.01, or < 0.001, 2-way ANOVA for panel A, Sidak post-test, Student’s t-test for other panels.

Treatment with fluorescently labeled siRNA (siRNA-AF-488) indicated that 50 nM siRNA was capable of entering human monocyte-derived myofibroblasts without the use of transfection reagents (Fig S1). The mixture of talin2 siRNA reduced not only the spindle-shaped morphology within the monocyte-derived myofibroblast population, but also the presence of FAs at the cell periphery, FBs within cells, and the localization of talin2 to the periphery of the cell (Fig S2).

In order to simplify treatment in anticipation of *in vivo* testing, we treated mouse myofibroblasts with individual siRNAs against mouse talin2, in addition to the same non-targeting, human talin2, and mouse talin2 siRNA mixtures as we treated the human monocytes. While each of the siRNA mixtures significantly decreased the number of αSMA and collagen I double-positive cells (Figure 4D), only talin2 siRNA #2 (sequence: 5’ CUGGAAAAUUCAGUGAUGA 3’ and antisense 5’ UCAUCACUGAAUUUUCCAG 3’) inhibited myofibroblast differentiation by itself. All mixtures and individual siRNAs decreased talin2 concentration within the cells (Figure 4E).

To determine if silencing talin2 can reduce fibroblast-myofibroblast differentiation, we added siRNAs individually and in mixtures at 50 nM to human and mouse fibroblast cultures. The mixture of human talin2 siRNA reduced both αSMA and collagen I double-positive (Fig 5A) and talin2 positive (Fig 5B) myofibroblasts. Mouse siRNA #2 most reduced both αSMA and collagen I double-positive (Fig 5C) and talin2 positive (Fig 5D) myofibroblasts in mouse cell cultures.

**Figure 5:**
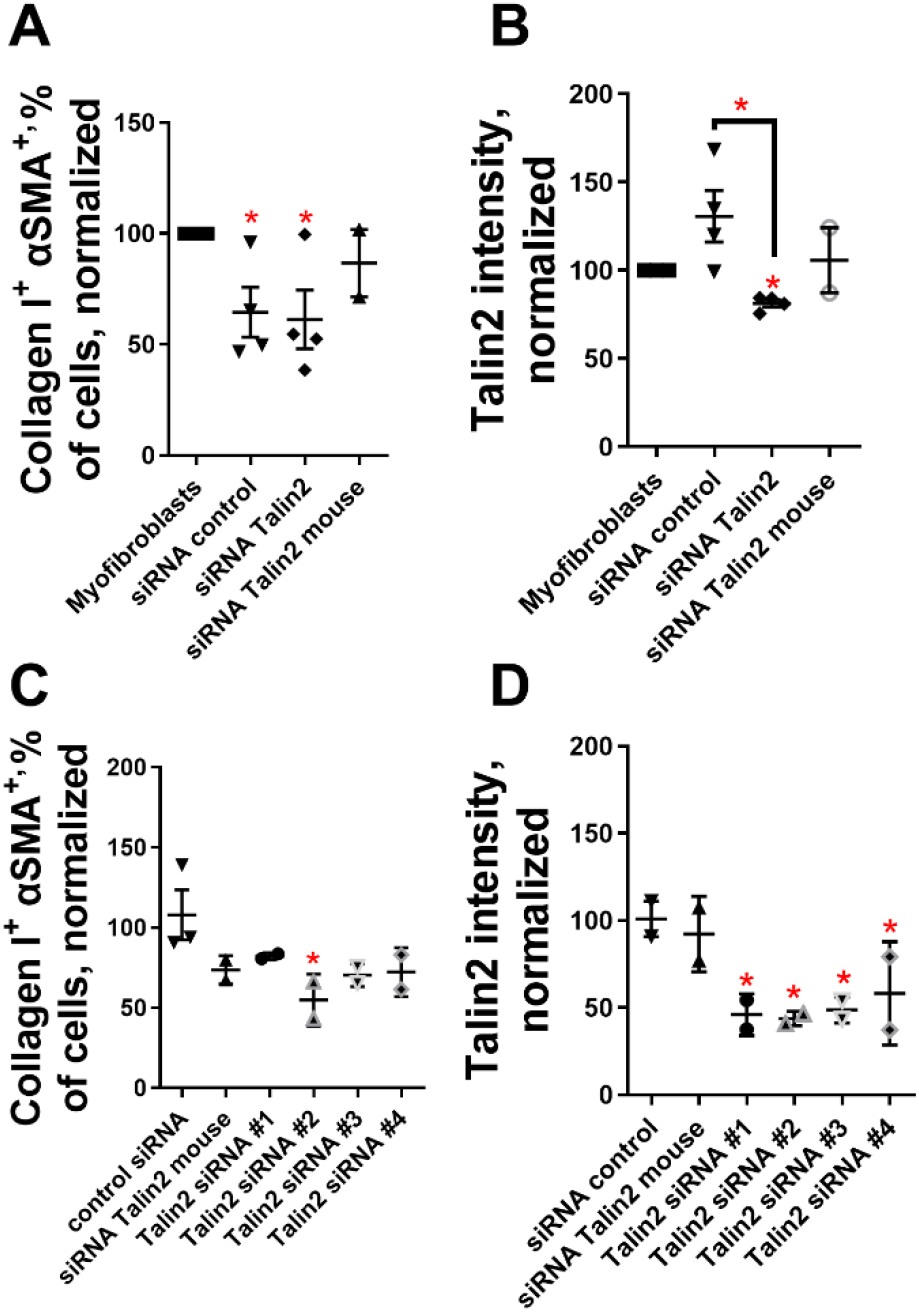
Silencing RNA against talin2 dedifferentiates human and mouse myofibroblasts. Fibroblasts were cultured on an infinite kPa tissue culture surface and treated with TGF-β, inducing them to become myofibroblasts. These myofibroblasts were treated with 50 nM mixtures of 4 control siRNAs, mixtures of 4 siRNAs targeting human talin2, mixtures of 4 siRNAs targeting mouse talin2, or 4 individual siRNAs targeting mouse talin2. Human myofibroblasts were assessed for (A) the number of αSMA^+^ Collagen I^+^ cells and (B) the amount of talin2, and mouse myofibroblasts were assessed (C) the number of αSMA^+^ Collagen I^+^ cells and (D) the amount of talin2. n ranges from 2 to 5. * = statistical significance of P < 0.05, < 0.01, or < 0.001, Student’s t-test.

To determine if reductions in myofibroblast’s spindle-shaped morphology, αSMA and collagen I content, and talin2 content correlated with a reduction in secreted pro-fibrotic factors, we assayed conditioned media from human and mouse monocyte-derived and fibroblast-derived myofibroblasts by ELISA. Treatment with talin2 siRNA did not significantly affect the amount of secreted anti-fibrotic IL-10 (Fig S3A) [48], but did reduce the amount of pro-fibrotic cytokines including IL-23 (Fig S3B) [49], CCL22 (Fig S3C) [50], IL-6 (Fig S3D) [51], CCL17 (Fig S3E) [50, 52], IL-12 subunit p40 (Fig S3F) [53], CXCL1 (Fig S3G and H) [54], TNF-α (Fig S3I) [55], and IL-1β (Fig S3J) [56]. While TGF-β was below the detection limit for this assay, IL-6 is sometimes sufficient to induce myofibroblast differentiation [57]. While the secretome for human cells was more frequently below the detection limit for this assay, similar results were seen, with reductions in pro-fibrotic TNF-α, MCP-1 [58], and IL-6 (Fig S4). This indicates that disruption of talin2 in not simply changing myofibroblast morphology, or talin2 localization, but is also dramatically altering the overall secretome of treated cells.

To determine if treatment with talin2 siRNA could rescue the damage from lung fibrosis in a treatment (not prophylactic) model, we insulted the lungs of male and female mice with bleomycin. We treated the mice via lung instillation with 200 nM of talin2 siRNA #2 at 7, 9, 11, 14, 16, and 18 days post insult, and euthanized the mice on day 21 post insult. While no treatment significantly altered mouse weight at day 21, treatment with talin2 siRNA showed a lesser transient weight decrease than did the controls (Fig 6A). Talin2 siRNA treatment also reduced the amount of collagen in the lungs as measured by a hydroxyproline assay (Fig 6B and C). Blinded Ashcroft scoring [59] of the Masson’s trichrome-stained lung sections, performed by another researcher, confirm that talin2 siRNA treatment improves the histological readout of lung fibrosis (Fig 6D). The individual talin2 siRNA treatment rescued lung fibrosis compared to non-targeting siRNA and untreated fibrotic lungs (Fig 6E-L). Thus, talin2 siRNA treatment rescues lung fibrosis in mice in both quantitative (hydroxyproline) and qualitative (Ashcroft scoring) measures, when treatment was provided 7 days after the inflammatory insult to the lung.

**Figure 6:**
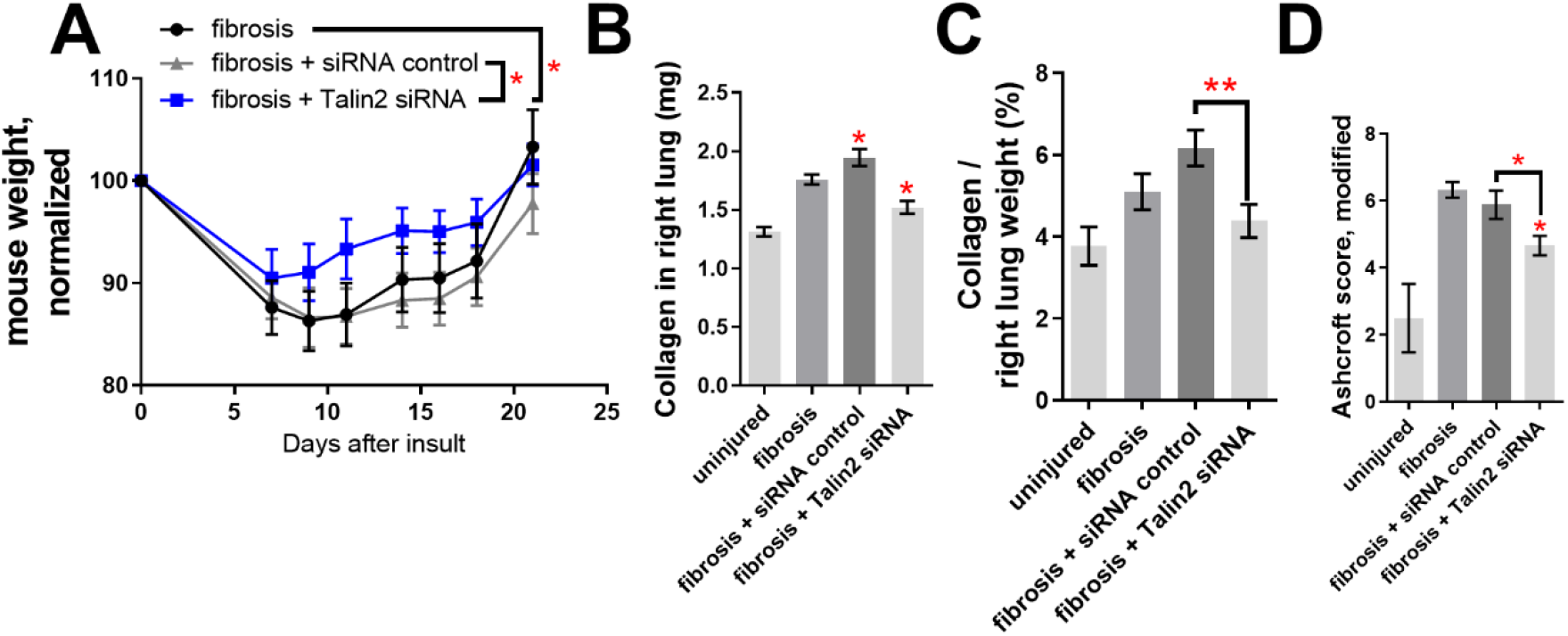
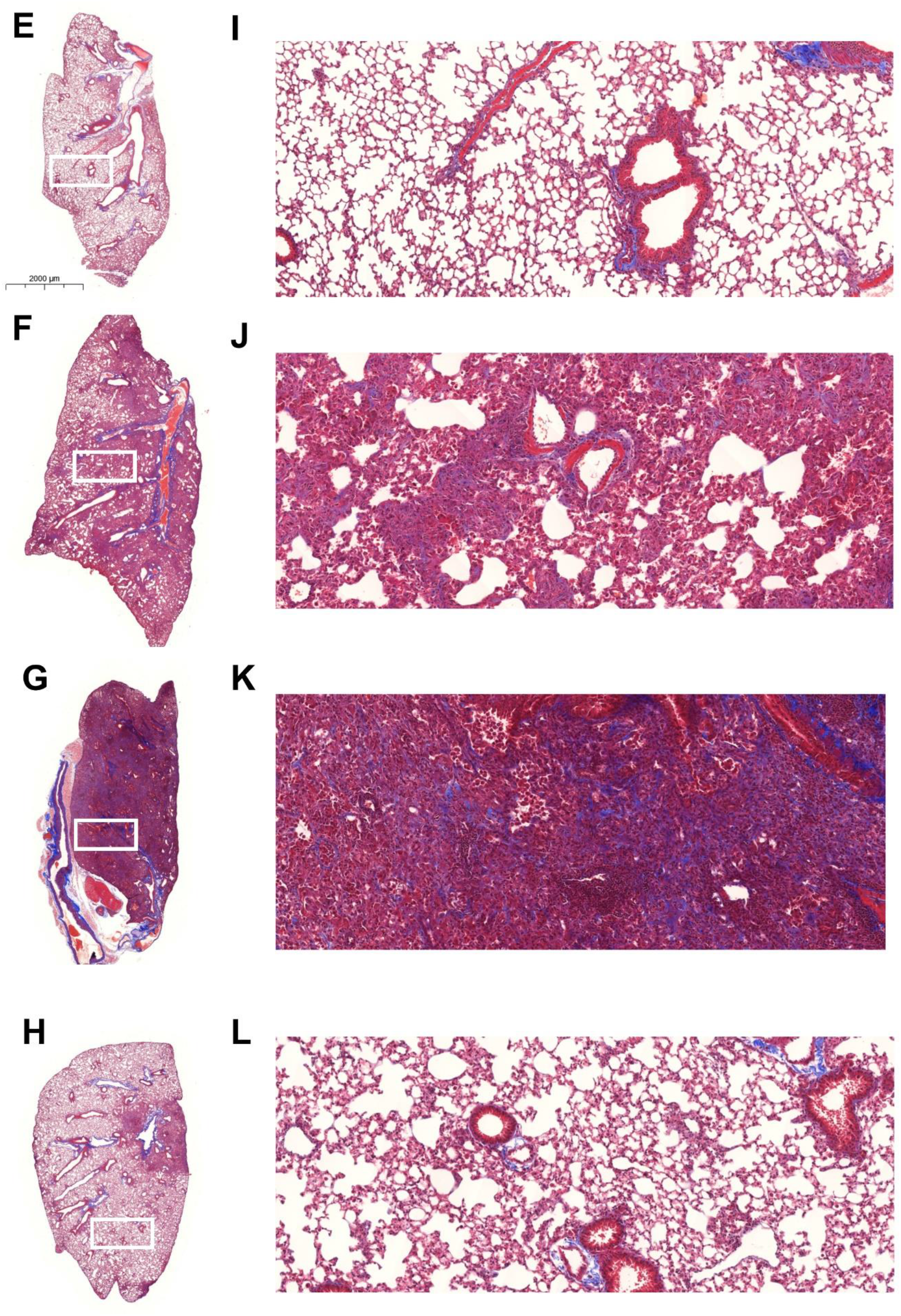
Talin2 siRNA rescues the fibrotic damage from bleomycin insult to mouse lungs. 50 μl of 0.2 μM talin2 siRNA was administered to mouse lungs 7, 9, 11, 14, 16, and 18 days after insult by bleomycin. (A) Mouse weights after treatment. (B) Collagen content from the right, multi-lobed lung assessed by hydroxyproline assay. (C) Data from C divided by dry weight of right lobes of mouse lungs. (D) Blinded Ashcroft scoring. (E-H) Representative images of left, single lobed lungs stained with Massons’s trichrome. (I-L) Inset of lungs. (E, I) Uninjured lung, (F, J) fibrotic lung, (G, K) fibrotic lung treated with siRNA control, and (H, L) fibrotic lung treated with talin2 siRNA. n ranges from 6 to 8. * = statistical significance of P < 0.05, < 0.01, or < 0.001, significance vs fibrotic lungs unless otherwise indicated, 2-way ANOVA for panel A, Sidak post-test, Student’s t-test for other panels.

To determine if treatment with talin2 siRNA improved lung fibrosis in both male and female mice, we analyzed the hydroxyproline-based collagen data (Fig S5A, B, D, E) and Ashcroft-based qualitative data (Fig S5C and F). The talin2 siRNA treatment improved collagen and Ashcroft readouts in male mice (Fig S5A-C), but only in the Ashcroft scoring for the female mice (Fig S5D-F), though this is only with n=3 mice per group.

Comparing the broncheo-alveolar lavage (BAL) of fibrotic and talin2 siRNA treated mouse lungs, there is reduced talin2 staining intensity in CD45^+^ cells, a reduced number of CD45^+^ talin2^+^ double positive cells, and a reduced number of CD45^+^ αSMA^+^ double positive cells (Fig S6).

To determine how well our talin2 siRNA treatment of fibrotic lungs compared with a complete reduction of talin2, we insulted the lungs of Tln2 -/- mice with bleomycin. Tln2 -/- lungs were resistant to fibrosis, showing reduced collagen deposition and a significantly improved Ashcroft score (Figure S7).

To determine if talin2 contributes to kidney fibrosis as well as lung fibrosis, we performed a unilateral ureteral obstruction (UUO) of the left kidney in Tln2-/- mice, allowed the kidneys to become fibrotic for 14 days, sacrificed the mice, and resected the kidneys. Tln2-/- mice are in the C57Bl6 background, so we used C57Bl6 mice as controls. The UUO-injured left C57Bl6 kidney (Figure 7B) showed significant fibrosis and damage, which was much reduced in the UUO-injured left Tln2-/- kidney (Figure 7D). The overall amount of collagen-I positive IHC stained tissue (brown) was much reduced in the Tln2-/- kidney (Figure 7E), and compared favorably to targeted anti-integrin treatments of kidney fibrosis [47]. Tln2-/- mice that have undergone UUO also have decreased blood urea nitrogen (BUN) and increased bilirubin levels compared to UUO-injured C57Bl6 mice (Figure S8). UUO-injured Tln2-/- mice also have higher alanine transferase (ALT) and aspartate transferase (AST) levels than UUO-injured C57Bl6 mice (Figure S8E and F). ALT and AST are commonly used markers of liver damage, but kidney fibrosis can counterintuitively cause ALT and AST levels to drop [60]. Several variables showed no significant change, including albumin, amylase, calcium, creatinine, creatine kinase, and uric acid.

**Figure 7:**
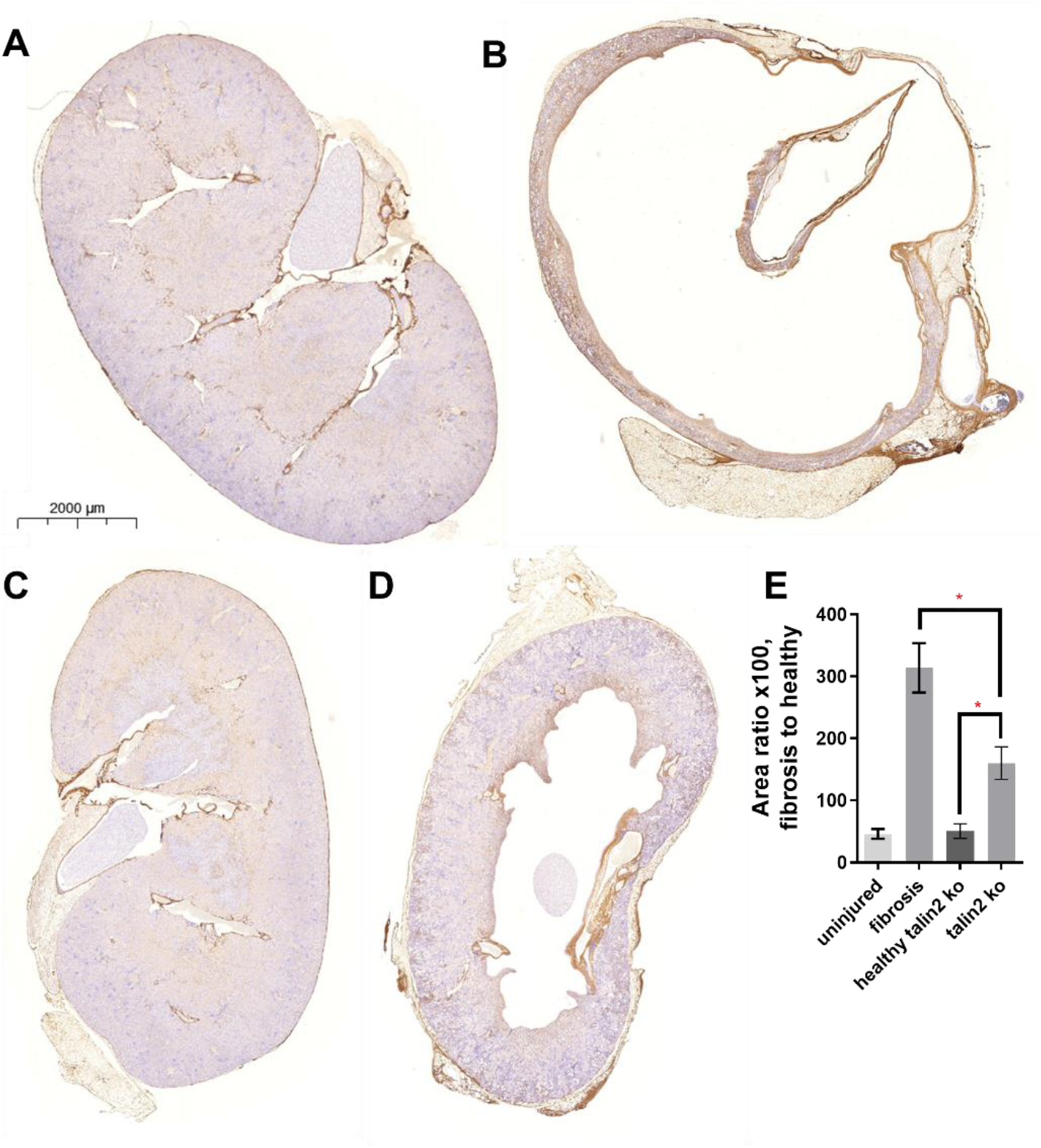
Tln2 -/- mice are resistant to UUO-induced kidney fibrosis. Mouse left kidneys were injured by UUO ligation. Kidneys were sectioned and IHC stained using α-collagen-I as a fibrosis marker. (A) Right, healthy C57Bl6 kidney. (B) Left, fibrotic C57Bl6 kidney. (C) Right, healthy Tln2 -/- kidney. (D) Left, fibrotic Tln2 -/- kidney. (E) The ratio of fibrotic to healthy tissue in each lung section. n = 6. * = statistical significance of P < 0.05, < 0.01, or < 0.001, Student’s t-test.

## DISCUSSION

In this paper, we show that adhesion mechanics plays a key role in myofibroblast differentiation (from monocytes and fibroblasts) and myofibroblast stability, and we show the utility of modulating adhesion machinery to propose a novel treatment for reversing fibrosis. We have shown that culture on somewhat stiffer (12 kPa) surfaces induces the expression of talin2, a key mechanosensing and mechanotransduction component of the cytoskeleton, compared to 1 kPa surfaces, which do not support monocyte-myofibroblast differentiation. Inhibition of talin2 can even de-differentiate myofibroblasts, changing their morphology, lowering the amount of αSMA and collagen I produced by myofibroblasts, and changing their secretion profile from pro- to anti-fibrotic. Inhibition of talin2 reverses established lung fibrosis in a murine model, and Tln2 -/- mice are resistant to bleomycin-induced lung fibrosis and UUO-induced kidney fibrosis.

Stepwise models of fibroblast-myofibroblast differentiation have previously been proposed that involve the presence of fibrosis-specific ECM and high extracellular stress [61]. Some studies have shown the importance of stiffness in the development of actin networks [62], finding that αSMA stress fibers begin to form at 3-6 kPa [63, 64], and form fully around 20 kPa [65, 66], consistent with our findings.

Fibroblast-myofibroblast differentiation on surfaces of different stiffness has also been investigated as it relates to morphology [35], and fibroblasts have been reported to form myofibroblasts beginning at 25 kPa [21]. Hepatocytes have also been shown to become myofibroblasts on a surface of 15 kPa [67], similar to our findings for monocytes and fibroblasts. Adhesion has also been recognized as important for monocyte differentiation [68], though not in the context of fibrosis.

Cells activate their mechanosensing machinery to assess the composition and stiffness of their environment, which allows them to respond to binding to a surface or a change in rigidity caused by tissue damage [62]. Classically, this activation begins when talin1 or talin2 activate integrins upon a change in their mechanical environment [30], the integrins then bind to the surface ECM. However, the order with which FAs come together has been called into question [69]. Monocytes could assess surface stiffness with talin1 or talin2 first, or assess the ECM composition of the surface using integrins. Previous studies have found pro-fibrotic [70, 71] and anti-fibrotic responses (Fig 6 in [72]) to various ECM coatings.

Talin1 and talin2 have overlapping but distinct (1) affinities for integrins and intracellular cytoskeletal proteins [73], (2) localizations within the cell [26], and (3) responsibilities in mechanosensing and mechanotransduction [74]. Because each talin isoform is mechanosensitive and part of a dynamic mechanosensing and mechanotransduction process, it can be difficult to universally parse what the responsibilities are for talins 1 and 2 [22]. The adhesome has been calculated to include almost 700 possible interactions [75]. Talin2 appears to be responsible for both the initial recruitment of FA and FB proteins at low elastic moduli, and also for mechanosensing and mechanotransduction through FBs, though not for mechanosensing in mature FAs [35]. However, knockout of talin2 has been shown to suppress large FA formation, while talin1 knockout allowed large FAs to remain [74]. Further complicating matters, talins also open multiple cryptic sites based on their response not just to surface stiffness, but also to tension, and fast and slow applications of force [22, 36].

Hinz proposes that super-mature FAs are a phenotypic checkpoint critical to a mechanical feedback loop of “extracellular stress and intracellular tension” that regulates the myofibroblast state [11]. Super-mature FAs form only under tension stress [65] and can be eliminated by re-culturing cells on softer surfaces [65, 66] or by reducing the FA’s maximum available size [11]. That super-mature FAs might be key control points matches well with our observation that adhesion mechanics modulates monocyte-myofibroblast differentiation. Perhaps the reason that monocytes cannot be induced to become myofibroblasts before being adhered or when adhered to soft surfaces is because they cannot form these super-mature FAs. Super-mature FAs may also be involved in a mechanism for scarless healing, as myofibroblasts that become stress-shielded by a fully repaired ECM may then lose the tension necessary to maintain super-mature FAs.

However, even with the diversity of talin1 and talin2 interactions with the cytoskeleton and adhesome [29], our immunofluorescence images also match phenotypes observed in other talin2 knockdown and knockout experiments. Hinz’s “super-mature focal adhesions” seen in fibroblasts look remarkably similar in size (30 um) and morphology to those seen in the monocyte-derived myofibroblasts shown in Figs S2A and S2B [25]. Similarly, Fig S2C appears phenotypically identical to monocytes cultured on a stiff surface but without pro-fibrotic signals to induce myofibroblast differentiation [73]. Interestingly, this similarity includes the subcellular localization of talin2 in macrophages [73]. The edges of talin2 siRNA-treated monocyte-myofibroblasts (Fig S2C) appear to have similarly diffuse actin staining, lacking FAs and FBs, similar to fibroblasts treated with talin2 siRNA [76] and in fibroblasts constitutively expressing a talin2 mutant that abrogated its interaction with the cytoskeleton [35]. This is not merely a difference in a macrophage vs myofibroblast phenotype, however, as cells from the monocyte-myofibroblast population that failed to form elongated, spindle shaped cells did display increased actin development and adhesion architecture (Fig S2D) compared to monocyte-myofibroblasts dedifferentiated with talin2 siRNA (Fig S2C).

Talin2 is key to both the formation of FAs [77] and FBs [26, 35] and is key to mechanotransduction and the generation of tension [35]. Talin1 and talin2 respond to cell stress and tension by unfolding in different ways, which exposes cryptic binding sites for integrins, actin, paxillin, and other members of the cytoskeleton and adhesome [33, 36]. Additionally, each subdomain of talin is itself mechanoresponsive and capable of binding to components of the members of the cytoskeleton and adhesome [33]. Reduction in the overall amount of talin2 might well be responsible for the reduction in super-mature FAs, which is turn could lead to the de-activation of myofibroblasts through apoptosis or de-differentiation [11].

Whether myofibroblasts remain active or de-activate determines whether damaged tissue repairs or scars, as repaired ECM is capable of stress-shielding resident cells and reducing the stress and tension on myofibroblasts [13, 78-80], which would prevent further scarring. Our goal was to induce myofibroblasts to de-activate (either through differentiation or apoptosis) by modulating adhesion machinery to convince myofibroblasts that they are no longer bound to a surface of sufficient stiffness. Several mechanisms have been proposed to de-activate myofibroblasts. De-differentiation has been achieved by re-culturing myofibroblasts on softer surfaces [65, 66, 81]. Interestingly, a recent investigation resulted in an experimental candidate drug that resolves fibrosis by means of apoptosis [20], based on changes in the metabolome of fibroblasts cultured on stiff surfaces.

Monocyte-myofibroblast differentiation was far more inhibited on low elastic modulus surfaces than was fibroblast-myofibroblast differentiation. This could be explained by a phenomenon called “mechanical-memory”, where fibroblasts that have been cultured long-term on functionally infinite modulus surfaces which are sufficient by themselves to induce myofibroblast differentiation [12], where the cells may carry with them a mechanical memory that makes de-differentiation difficult to achieve over short timescales [21].

Talin2 is an excellent target for disrupting a myofibroblast’s mechanosensing capability that determines whether a myofibroblast remains a myofibroblast. Talin2 provides a framework on which all other components of the adhesome and cytoskeleton [33], and is involved in mechanosensing [35, 77] and mechanotransduction [35]. Knockouts of talin2 have few (or no) phenotypic irregularities [28, 38], while talin1 knockouts are embryonic lethal [37]. Talin1 is expressed ubiquitously, while talin2 is expressed on monocyte-derived cells, in muscle tissue, and in the brain [28, 31, 32]. Talin2 transcripts are also upregulated in monocyte-derived cells in idiopathic pulmonary fibrosis (IPF, Fig S9, data from the IPF cell atlas [82]). Thus, in seeking to disrupt mechanosensing in monocytes, we turned our attention to talin2 rather than talin1.

To our knowledge, the only method of targeting talin other than siRNA is the small molecule KCH-1521, which decreases adhesion-dependent angiogenesis. KCH-1521, however, does not distinguish between talin1 and talin2 [83]. By contrast, siRNA treatments permit a distinction between talin2 and talin1. Fortunately, Accell siRNA offers promise for delivering talin2 knockdown to the lungs, and it could potentially even be delivered as a dry powder by inhalation [84]. Our doses are a factor of 10- to 100-fold lower than have been used in a previous pulmonary study using Accell siRNA [85].

## MATERIALS AND METHODS

### Study Design

This study examines two related hypotheses. The first is that myofibroblast differentiation (from monocytes and fibroblasts) is governed by adhesion mechanics, and the second is that modulation of adhesion mechanics machinery can be used to guide novel treatment discovery for fibrosing diseases. Specifically, we tested whether inhibition of the intracellular tension sensor talin2 could be used to treat lung fibrosis in mice. Group size was selected based on experience with the pulmonary fibrosis model, particularly based on a pilot experiment using talin2 knockdown in lung fibrosis. Mice were randomized into treatment groups within a cage to eliminate cage effects from the experiment. Treatment was performed by multiple researchers over the course of this study, to ensure reproducibility. Lungs were also resected by multiple researchers, and blinded scoring was used on the fibrosis histology images.

### Purification of human monocytes

In order to isolate more peripheral blood mononuclear cells (PBMC) than is possible through a single blood donation, PBMCs were purified from leukocyte reduction filters obtained from the University of Chicago blood donation center, in accordance with human subject protocol at the University of Chicago. All leukocyte reduction filters were de-identified and taken from random blood donors regardless of age, race, or gender. Blood was filtered, and leukocytes purified, the same afternoon as a morning blood donation, to reduce the amount of time that PBMCs could adhere to the filter.

The leukocyte reduction filter was sterilized with 70% ethanol, and the blood flow tubes on either end were clamped to prevent flow. The tube through which filtered blood had exited the filter was cut below the clamp, and a syringe containing 60 ml phosphate buffered saline (PBS) was inserted into the tube. Following this, the tube through which unfiltered blood had entered the filter was unclamped, and PBS was slowly pushed through the filter in the opposite direction of the original blood flow. This direction of flow provided the highest recovery of PBMC, approximately 300 million cells per filter.

The collected blood from the filter was then layered with lymphocyte separation media (LSM), and centrifuged at 1300 xG for 20 min. The PBMC layer was then removed by pipetting.

These PBMCs were further purified by use of a negative selection kit for human monocytes (Stemcell, Cambridge, MA), per the manufacturer’s instructions. The expected yield from each filter was approximately 20 million monocytes. Monocytes were then washed by PBS using five successive 300 xG centrifugation steps, in order to remove EDTA from the resulting population. Monocytes were checked for purity using flow cytometry, and average purity was above 95%. Monocytes were cultured immediately following purification, at 100,000 monocytes/cm^2^.

### Purification of mouse monocytes

All PBS, plasticware, filters, glassware and magnets were pre-chilled to 4C, and kept cold throughout this procedure, to limit the clumping of cells. Importantly, ACK lysis buffer caused cell death, and so was not used in this procedure.

Spleens were resected from healthy C57BL/6 mice, pooled, and were placed in PBS with 1 mM EDTA, 2% fetal calf serum (FCS) to prevent subsequent clumping of cells. Spleens were pushed through a 100 mm filter to disassociate the cells. Monocytes were purified from disassociated cells by use of a negative selection kit (Stemcell), following the manufacturer’s instructions. The purified monocytes were then washed by PBS using five successive 300 xG centrifugation steps, in order to remove EDTA from the resulting population. Monocytes were checked for purity using flow cytometry, and average purity was above 95%. The average yield was 1.5 million monocytes per spleen.

Monocytes were cultured immediately following purification, at 250,000 monocytes/cm^2^.

### Culture of human and mouse monocytes

Human and mouse monocytes were cultured as previously described, using serum-free media (SFM) [86]. Briefly, SFM for human cells is composed of fibrolife (Lifeline, Frederick, MD), with 1x ITS-3 (Sigma, St. Louis, MO), 1x HEPES buffer (Sigma), 1x non-essential amino acids (Sigma), 1x sodium pyruvate (Sigma), and penicillin-streptomycin with glutamate (Sigma). For mice, 2x concentrations of ITS-3, HEPES buffer, non-essential amino acids, and sodium pyruvate are added. For mouse monocytes, 50 mM beta-mercaptoenthanol (ThermoFisher) was also added, as were pro-fibrotic supplements M-CSF (25 ng/ml, Peprotech, Rocky Hill, NJ) and IL-13 (50 ng/ml). Additionally, M-CSF and IL-13 were refreshed in the media of mouse monocytes after 3 days of culture.

Addition of tryptase (Fitzgerald, Acton, MA) to media was as previously described [87]. Monocytes were allowed to differentiate for 5 days, and counted based on morphology as previously described [88].

### Culture of human and mouse fibroblasts

Human fibroblasts (MRC-5, ATCC, Manassas, VA) and mouse fibroblasts (NIH-3T3, ATCC) were cultured in SFM composed for human cells, with 1x concentrations of additives. TGF-b (Peprotech) was added to induce myofibroblast formation at 5 ng/ml [89]. Cells were cultured at 10,000/cm^2^, and TGF-β was refreshed in cultures weekly to maintain the myofibroblast phenotype.

### Methods for rolling monocytes in the presence of pro-fibrotic factors

100,000 human monocytes were suspended in SFM, and placed onto tissue culture treated plastic, or into low adhesion microcentrifuge tubes (protein lo-bind, ThermoFisher). Tryptase was added at 12.5 ng/ml [87], while IL-13 was added at 50 ng/ml. Monocytes were incubated for 1 hr, either adhered to the tissue-culture treated plastic or rotating in low adhesion microcentrifuge tubes. After 1 hr, the adhered monocytes and rotating monocytes were gently washed with four successive PBS washes to remove both tryptase and IL-13 from the surface of the monocytes. The washed monocytes (including those from low adhesion microcentrifuge tubes) were then cultured on tissue culture treated plastic for 5 days, and the number of myofibroblasts was observed through counting of spindle shaped morphological cells and through analysis of alpha-smooth muscle actin positive (αSMA^+^) and collagen I ^+^ cells. αSMA and collagen I are widely used markers of myofibroblast differentiation [90].

### Preparation of low surface stiffness plates

Low stiffness tissue-culture plates were ordered from Matrigen (San Diego, CA). Culture of myofibroblasts on low stiffness surfaces was similar to the culture conditions on tissue-culture treated plasticware. However, low stiffness surfaces were coated in 10 mg/ml fibronectin (Millipore Sigma) for 1 hr at 37C. Unbound fibronectin was gently removed with 3 successive PBS changes. For experiments involving the comparison of infinite binding surfaces to softer surfaces, the tissue culture treated plasticware was also fibronectin coated.

### mRNA purification and RNAseq

mRNA was purified using the Trizol method [91]. RNA sequencing was performed by the University of Chicago Center for Research Informatics, and full protocols for RNA sequencing are in supplementary File 1.

### siRNA treatment of human and mouse cells

Monocytes are difficult to transfect [92]. As monocytes are non-proliferating cells [92], plasmid-based siRNA platforms that require entry into the nucleus are also unlikely to inhibit protein production across the entire population.

Accell Silencing RNA (siRNA, Dharmacon, Lafayette, CO) is specifically engineered to penetrate cells without the use of transfection reagents, due to a cholesterol modification at one terminus of the sequence. Additionally, Accell siRNA is well suited for a study for silencing proteins in otherwise non-proliferating cells, because each non-replicating siRNA molecule does not have to enter the nucleus, a necessary prerequisite for plasmid-based siRNA platforms. Thus, Accell siRNA targets all cells in a population, not just those that are replicating. Further, that the Accell siRNA does not replicate inside a cell is ideal from a dose-recovery standpoint. Lastly, Accell siRNA contains a modification that inhibits enzymatic digestion within cells. Thus, Accell siRNA made a natural choice for this experiment, and all siRNAs used in this study are Accell siRNA. siRNA was resuspended in RNAse-free ultrapure water, as per the manufacturer’s instructions.

The SMARTpool of siRNA targeting human talin2 contained four target sequences, while the anti-parallel corresponding siRNA sequences actually silenced the mRNA. Target sequences: #1 (CCAGAAAACUGAACGAUUA), #2 (GCCCUGUCCUUAAAGAUUU), #3 (CGACUGUGGUUAAAUACUC), and #4 (CGAGAAAGCUUGUGAGUUU). The SMARTpool targeting mouse talin2 also contained four target sequences: #1 (CGACUGUGGUUAAAUACUC), #2 (CUGGAAAAUUCAGUGAUGA), #3 (CCCUGGAUUUUGAAGAACA), and #4 (CCAUCGAGUACAUAAAACA). Control non-targeting SMARTpools contained 4 target sequences, including: #1 (CCAGAAAACUGAACGAUUA), #2 (GCCCUGUCCUUAAAGAUUU), #3 (CGACUGUGGUUAAAUACUC), and #4 (CGAGAAAGCUUGUGAGUUU). Individual siRNA tested in mice was talin2 siRNA #2, target: 5’ CUGGAAAAUUCAGUGAUGA 3’, anti-sense: 5’ UCAUCACUGAAUUUUCCAG 3’. Control non-targeting siRNA used in the mouse in vivo study was (CCAGAAAACUGAACGAUUA).

### Flow cytometry

Myofibroblasts were removed from their tissue culture surfaces by the use of cold trypsin-EDTA (Sigma), followed by mechanical agitation by a rubber policeman. Cells were fixed and permeabilized using Cytofix/Cytoperm (BD biosciences, Franklin Lakes, NJ). Live dead stain was live-dead aqua, used per manufacturer’s instructions (ThermoFisher). Compensation was performed via UltraComp beads (ThermoFisher) per the manufacturer’s instructions. Antibodies used were anti-collagen I (Biolegend, San Diego, CA), anti-alpha smooth muscle actin (αSMA) (R and D systems, Minneapolis, MN), anti ki-67 (BD biosciences), and anti-talin2 (R and D systems).

### siRNA penetration into cells

siRNA complexed to AF-488 (Dharmacon) was added to human and mouse monocytes and fibroblasts at 50 nM, to determine if the siRNA was penetrating the cell and remaining for up to 2 days. Cells were removed from their culture surface as previously indicated, fixed, and analyzed via flow cytometry.

### Mouse lung instillation

All mouse experiments were performed under supervision with protocols approved by the University of Chicago IACUC. Male and female mice were acquired at 8 weeks of age (Jackson laboratories, Bar Harbor, ME) with the intent to be used at 12 weeks of age. However, due to delays regarding COVID19 lockdown and the allowed resumption of non-COVID19 research, the mice were 32 weeks old when the study progressed. Mouse lungs were instilled with bleomycin (0.075 units, 75 ug, Fresenius Kabi, Switzerland) and siRNA (0.2 uM), suspended in endotoxin-free PBS, as previously described [93]. First, mice were anesthetized via isoflurane inhalation (2%). Mice were then placed upright on an angled surface, their tongue pulled to the side, and a 200 ml narrow pipet was placed at the entrance of their throat. 50 ml of PBS was dispensed to the entrance of the throat, and mice were allowed to inhale. Administration to the lungs was confirmed by listening to the mouse’s breathing for popping noises. Mice were then weighed and placed on a heating pad to recuperate.

Following bleomycin insult, mice were treated with siRNA using an identical installation procedure. The dose schedule was 7, 9, 11, 14, 16, and 18 days following bleomycin insult. Mice were euthanized at 21 days post insult via injecting of euthasol (Covetrus, Portland, ME) instead of CO_2_ inhalation, which could damage the lungs.

### Lung resection and fibrosis scoring

Lungs were resected, and perfused with 5 ml of PBS via cardiac puncture. BAL involved exposing the trachea and penetrating the trachea using an 18 gauge needle. The BAL was performed by inserting a catheter needle (Exel International, Redondo Beach, CA) into the trachea, and slowly moving 800 ml of PBS into and out of the lungs. If blood entered the lavage it was discarded. In a previous pilot study testing the efficacy of talin2 siRNA treatment, the lungs were then broncheo-alveolar lavaged (BAL), and the resulting lavage frozen in 10% DMSO. Due to limitations associated with COVID19 scheduling, BAL was not possible in the larger study. This lavage was thawed, fixed, immunostained, and analyzed via flow cytometry to provide the data on talin2’s in vivo reduction following siRNA treatment.

After resection, the right and left lobes were separated. The left lobe was fixed in 4% paraformaldehyde overnight, mounted in paraffin, sectioned into 5 mm slices, and stained using Masson’s trichrome. Stained lungs were scanned at high resolution using a CRi Panoramic SCAN 40x Whole Slide Scanner (Perkin-Elmer, Waltham, MA), and were read for fibrosis using a modified Ashcroft method, as previously described [59].

The right lobe of the lung was frozen, and dehydrated using a tissue lyophilizer (Labconco, Kansas City, MO). This dehydrated lung was weighed, and was assessed for collagen content by hydroxyproline assay, as previously described [94]. Briefly, dried lungs were digested in 6N HCl/PBS at 100C for 24 hr. The supernatant from this digestion was added to 96 well plates and treated sequentially with chloramine-T solution and Ehrlich’s solution at 65C for 15 min to facilitate the color change reaction. Color was read at 561 nm. Quantification was provided by use of a hydroxyproline (Sigma) dilution series, which was transformed into a standard curve.

### Tln2-/- mice

Tln2-/- mice (MGI 1917799) were the kind gift of Drs. Roy Zent and David Critchley. Mouse genomic DNA was isolated through ear punches, and Tln2-/- mice were validated genomic DNA by PCR (Transnetyx).

### Kidney unilateral ureteral obstruction (UUO) fibrosis model

UUO surgery was performed as previously described [95], with adjustments. Briefly, Mice were anesthetized via 2% isoflurane inhalation, and injected with meloxicam (1 mg/kg), buprenorphine (0.1 mg/kg) in a saline solution, subcutaneously. Briefly, mice were laid on their right side and an abdominal incision used to visualize the left ureter. The left ureter was ligated in the middle section of the ureter with two ties (2mm apart) using 7-0 silk sutures. Peritoneum is then closed with 5-0 vicryl and skin is closed with 5-0 nylon. 14 days following the UUO ligation, the mice were sacrificed and the kidneys resected. In each case, the UUO ligation was still in place, and in each case it was.

### Assessment of fibrosis in kidneys

Right and left kidneys were placed in 4% PFA for 24 hours, mounted in paraffin, sectioned into 5 mm full kidney slices, and stained using immunohistochemistry for collagen I (1:4000, polyclonal rabbit, lifespan biosciences, Seattle WA) via a Bond-Max autostaining system (Leica biosystems, Lincolnshire, IL). Stained kidneys were scanned at high resolution using a CRi Panoramic SCAN 40x Whole Slide Scanner (Perkin-Elmer).

Images were equalized in size, and converted to .tif files using CaseViewer. Images were then imported into imageJ, scale set for conversion between microns and pixels, and deconvoluted with the “H DAB” deconvolution option. The blue stain was thresholded at 215 to see how many pixels were positive for nuclear staining but negative for collagen I, and the brown (IHC positive) image thresholded at 185 to see how many pixels were positive for collagen I.

Machine-staining allowed these kidneys to be compared with high reproducibility.

### Blood analysis for markers of kidney damage

At the time of euthanasia, blood was collected via submandibular bleed into protein low-bind tubes and allowed to coagulate for 2 hours on ice. Coagulated blood was then centrifuged at 10,000 xG for 10 min, and serum collected. Serum was then diluted 4x in MilliQ water before being placed on deck on an Alfa Wassermann VetAxcel Blood Chemistry Analyzer. All tests requiring calibration were calibrated on the day of analysis and quality controls were run before analyzing samples. Serum tests were run according to kit instructions, and Creatinine Kinase was normalized to calcium ion concentrations where indicated to account for sample hemolysis.

### Immunofluorescence

Human monocytes from 3 donors were cultured in 8-well chamber slides (Millipore-Sigma) in SFM, and allowed to become myofibroblasts over 3 days. Myofibroblasts were then treated with SMARTpools for non-targeting control siRNA and siRNA targeting human talin2. After 1 week of de-differentiation, the slides were dried quickly using the airflow from a laminar flow hood, in order to preserve cellular morphology as accurately as possible. Cells were then fixed with ice cold 4% PFA, permeabilized with saponin (Sigma). Primary antibodies (anti-talin2, novus) were added at 5 mg/ml overnight. Cells were washed 3x in PBS, and were exposed to DAPI and F-actin-phalloidin-488 (ThermoFisher) for 1 min. Cells were mounted using water-based mowiol mounting media (Southern Biotech, Birmingham, AL) to preserve fluorescence. Slides were imaged immediately using a confocal microscope (Olympus, Shinjuku City, Tokyo).

### Legendplex ELISA

Supernatant from experiments involving culturing human and mouse monocytes and fibroblasts into myofibroblasts on surfaces of different stiffnesses and treatment with siRNA were thawed at 4C, and centrifuged at 4c and 10,000 xG force to pellet cell debris. Supernatant was taken and added to plates, in triplicate. Legendplex (Biolegend) beads against general inflammation markers were added, according to the manufacturer’s instructions. Sample readouts were normalized to each individual experiment and donor control.

### Statistical analysis

Statistical analyses were performed using GraphPad Prism software, and P < 0.05 was considered statistically significant. 2-way ANOVA and Student’s t-test were used to compare groups.

## Acknowledgements

We thank the Human Tissue Resource Center of the University of Chicago for histology analysis. We thank the Integrated Light Microscopy Core of the University of Chicago for Imaging. We thank the Genomics core at the University of Chicago for RNAseq analysis. We’d also like to acknowledge blind scoring from Margo MacDonald, and guidance on fibrosis models from Anne Sperling.

## Abbreviations

(αSMA): alpha-smooth muscle actin
(ALT): Alanine Transferase
(AST): Aspartate Transferase
(BAL): broncheo-alveolar lavaged
(BUN): blood urea nitrogen
(FCS): fetal calf serum
(FAs): focal adhesions
(FBs): Fibrillar adhesions
(IPF): idiopathic pulmonary fibrosis
(IHC): Immunohistochemistry
(MRC-5): Human fibroblasts
(kilopascals): kPa
(MCP1): macrophage-chemotactic protein-1
(NIH-3T3): mouse fibroblasts
(PBMC): peripheral blood mononuclear cells
(PBS): phosphate buffered saline
(SFM): Silencing RNA
(siRNA): serum-free media
(TGFβ): Transforming growth factor β
(Tln2): Talin2
(TNFα): Tumor necrosis factor α
(UUO): Unilateral Ureteral Obstruction

## Funding

This work was supported in part by the University of Chicago (to JAH).

## Author’s contributions

Conceptualization: MJVW

Methodology: MJVW

Investigation: MJVW, MO, JEGM

Visualization: MJVW

Funding acquisition: JAH

Project administration: MJVW, JAH Supervision: MJVW, JAH

Writing – original draft: MJVW

Writing – review & editing: MJVW, JAH

## Ethics approval

All the animal experiments performed in this work were approved by the Institutional Animal Care and Use Committee of the University of Chicago.

## Conflicts of interest

MJVW and JAH are inventors on U.S. Patent Application No. 63/196,594. The other authors declare that they have no competing interests.

## Tables

**Supplemental Table 1.**
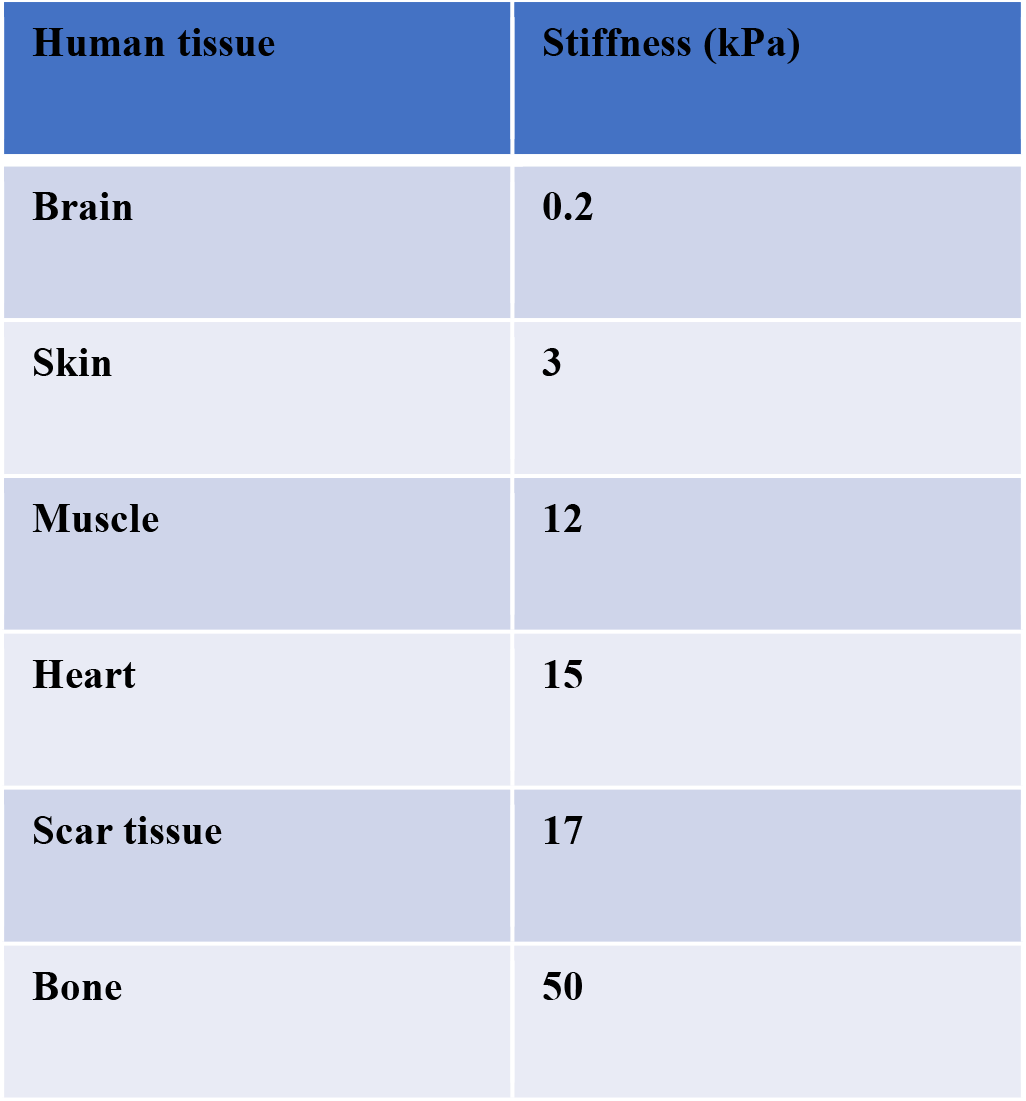
Representative human tissue measurements for Young’s elastic modulus.

**Supplemental table 2.**
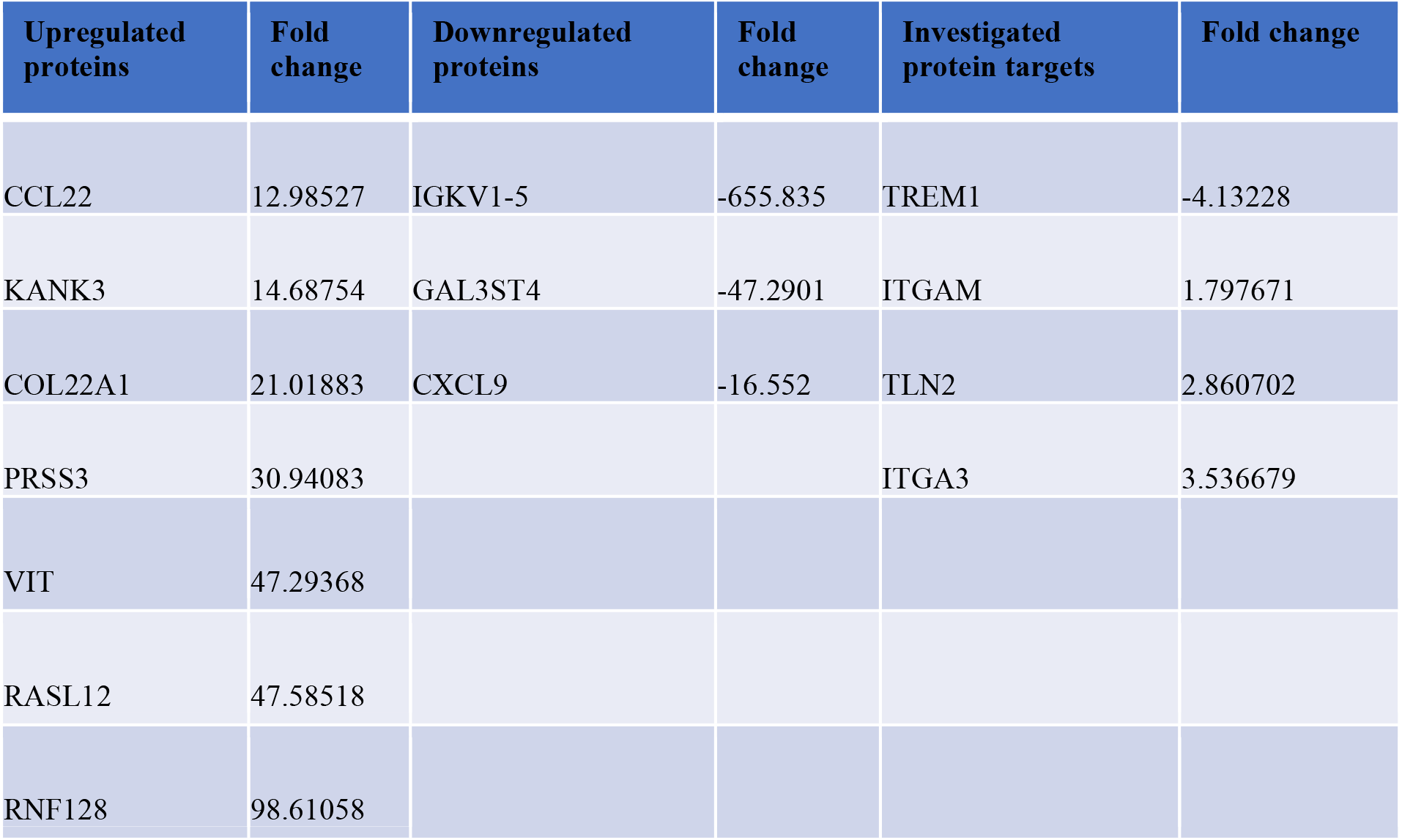
Upregulated and downregulated RNAseq transcripts.

| Upregulated proteins | Fold change | Downregulated proteins | Fold change | Investigated protein targets | Fold change |
| --- | --- | --- | --- | --- | --- |
| CCL22 | 12.98527 | IGKV1-5 | -655.835 | TREM1 | -4.13228 |
| KANK3 | 14.68754 | GAL3ST4 | -47.2901 | ITGAM | 1.797671 |
| COL22A1 | 21.01883 | CXCL9 | -16.552 | TLN2 | 2.860702 |
| PRSS3 | 30.94083 |  |  | ITGA3 | 3.536679 |
| VIT | 47.29368 |  |  |  |  |
| RASL12 | 47.58518 |  |  |  |  |
| RNF128 | 98.61058 |  |  |  |  |

**Supplemental table 3.** Upregulated and downregulated RNAseq pathways, and functional annotation clustering. Enrichment score is calculated based on the maximum deviation from 0 in the pathway analysis [96]. In this analysis, 5.18 is the highest possible score.

| Upregulated pathways | Downregulated pathways | Functional annotation clustering | Enrichment score |
| --- | --- | --- | --- |
| Chemokine signaling | Role of nfat in regulation of the immune response | Complement activation | 5.18 |
| Leukocyte extravasation signaling | TREM1 | Chemokine signaling | 5.18 |
| IL-6 signaling | PTEN signaling | Phagocytosis recognition | 5.18 |
| Paxillin signaling | Coagulation system | positive regulation of inflammatory response | 5.18 |
| Integrin signaling |  | monocyte chemotaxis | 5.18 |
| IL-13 signaling |  | receptor-mediated endocytosis | 5.18 |

## Supplementary Materials

**Supplemental figure 1:**
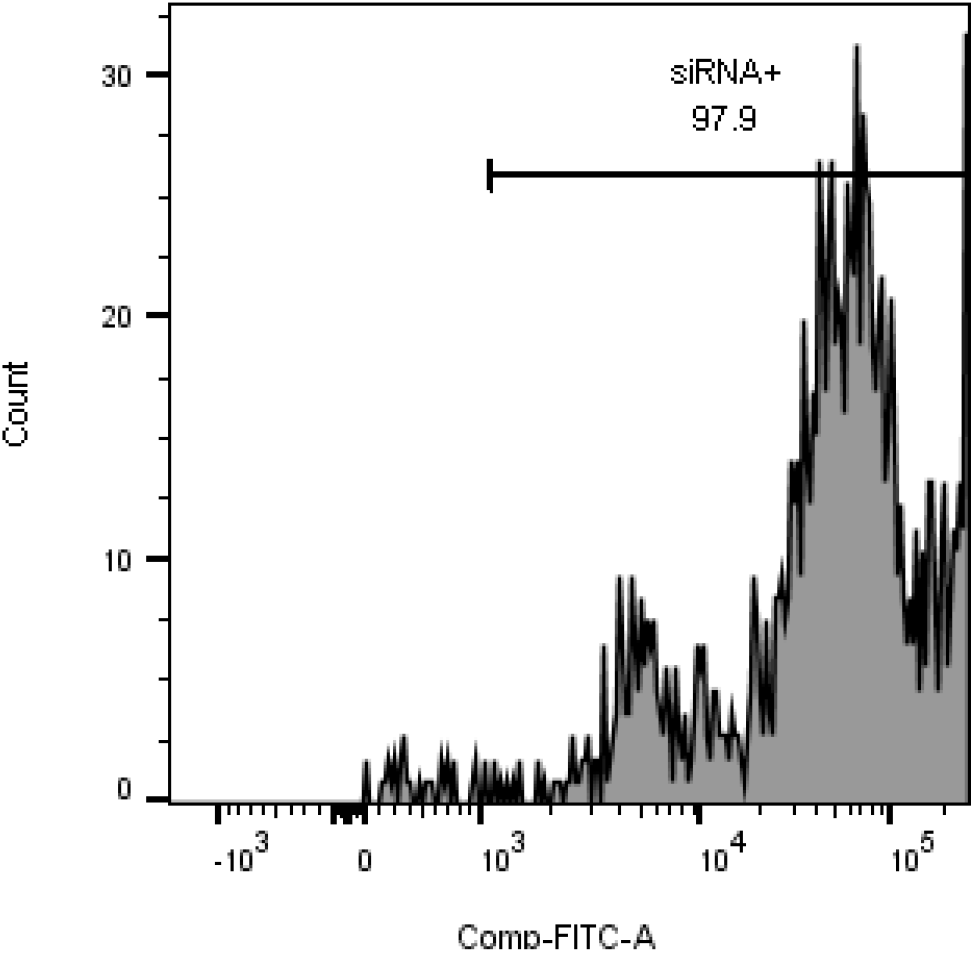
The addition of siRNA-AF-488 to human monocyte-myofibroblasts shows an increase in the amount of green fluorescence in cells.

**Supplemental figure 2:**
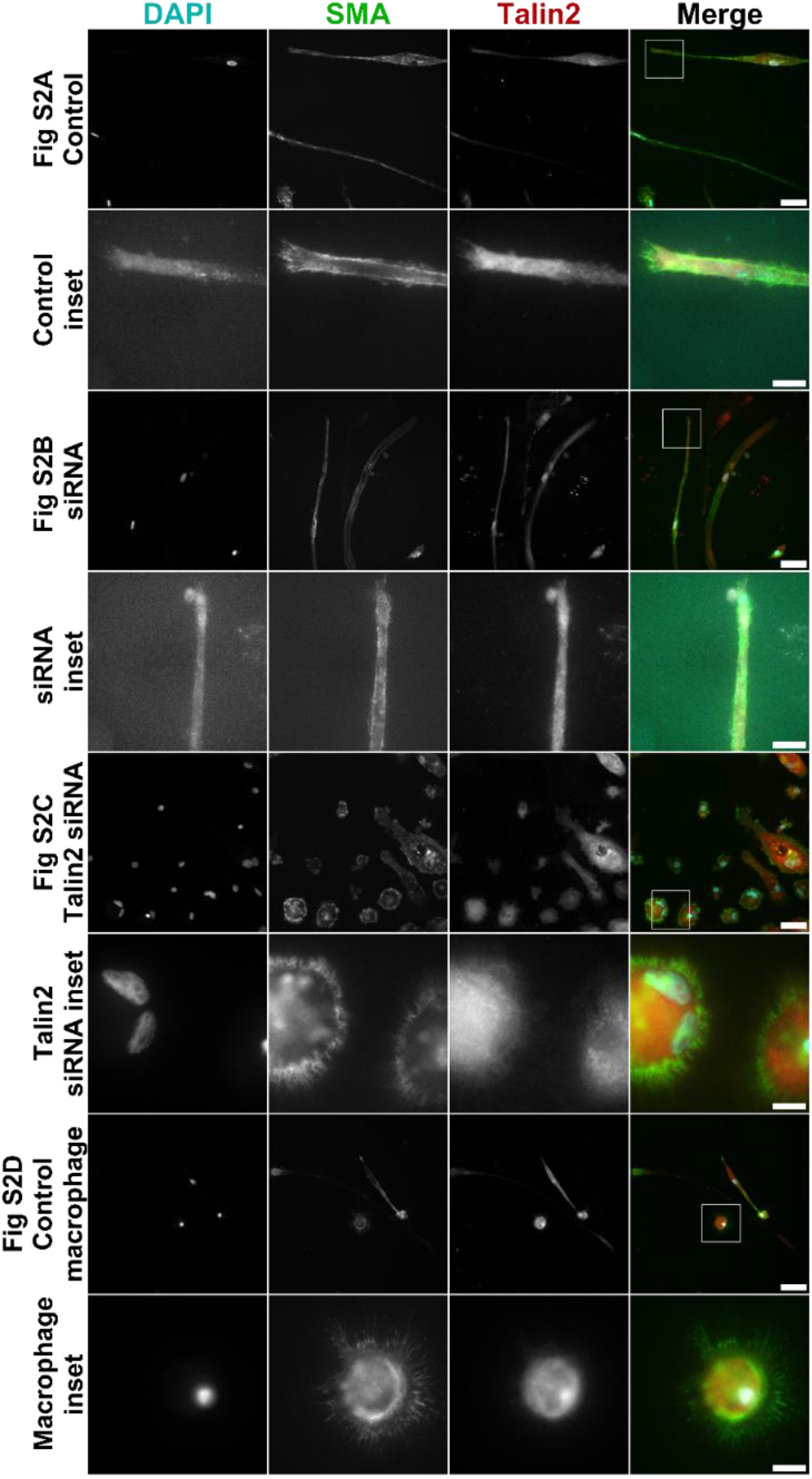
Treatment with talin2 siRNA alters the morphology of myofibroblasts, as well as the localization of talin2. Human monocyte-derived myofibroblasts were treated with 50 nM siRNA targeting talin2. (A) A myofibroblast representative of morphology. (B) Untreated population of myofibroblasts, inset on nucleus and focal adhesion at the tip of the myofibroblast. (C) Untreated population of myofibroblasts, clear talin2 localization at the periphery of the cell. (D) Control siRNA treated myofibroblasts, inset on tip of myofibroblast. (E) Talin2 siRNA treated myofibroblasts, inset on edges of de-differentiated myofibroblasts. Red = talin2, green = αSMA, blue = DAPI. Size bar = 30 μm, inset size bar = 10 μm.

**Supplemental figure 3:**
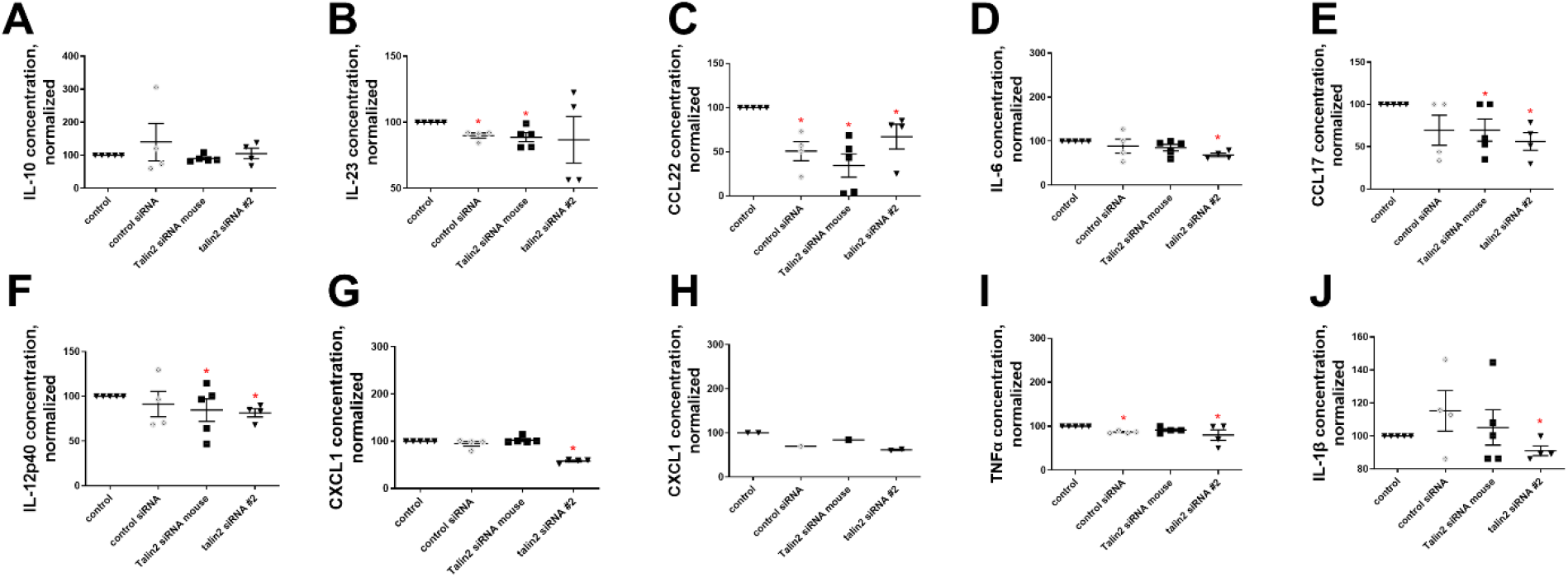
Treatment with talin2 siRNA reduces the pro-fibrotic secretome of mouse myofibroblasts. Treatment with 50 nM of talin2 siRNA on mouse monocyte-derived myofibroblasts does not affect the amount of secreted (A) IL-10, but reduces the amount of secreted (B) IL-23, (C) CCL22, (D) IL-6, (E) CCL17, (F) IL-12 subunit p40, (G) CXCL1, (I)TNF-a, and (J) IL-1β. (H) is from the secretome of human fibroblast-myofibroblasts. n ranges from 3 to 4. * = statistical significance of P < 0.05, < 0.01, or < 0.001, significance vs control unless otherwise indicated, Student’s t-test.

**Supplemental figure 4:**
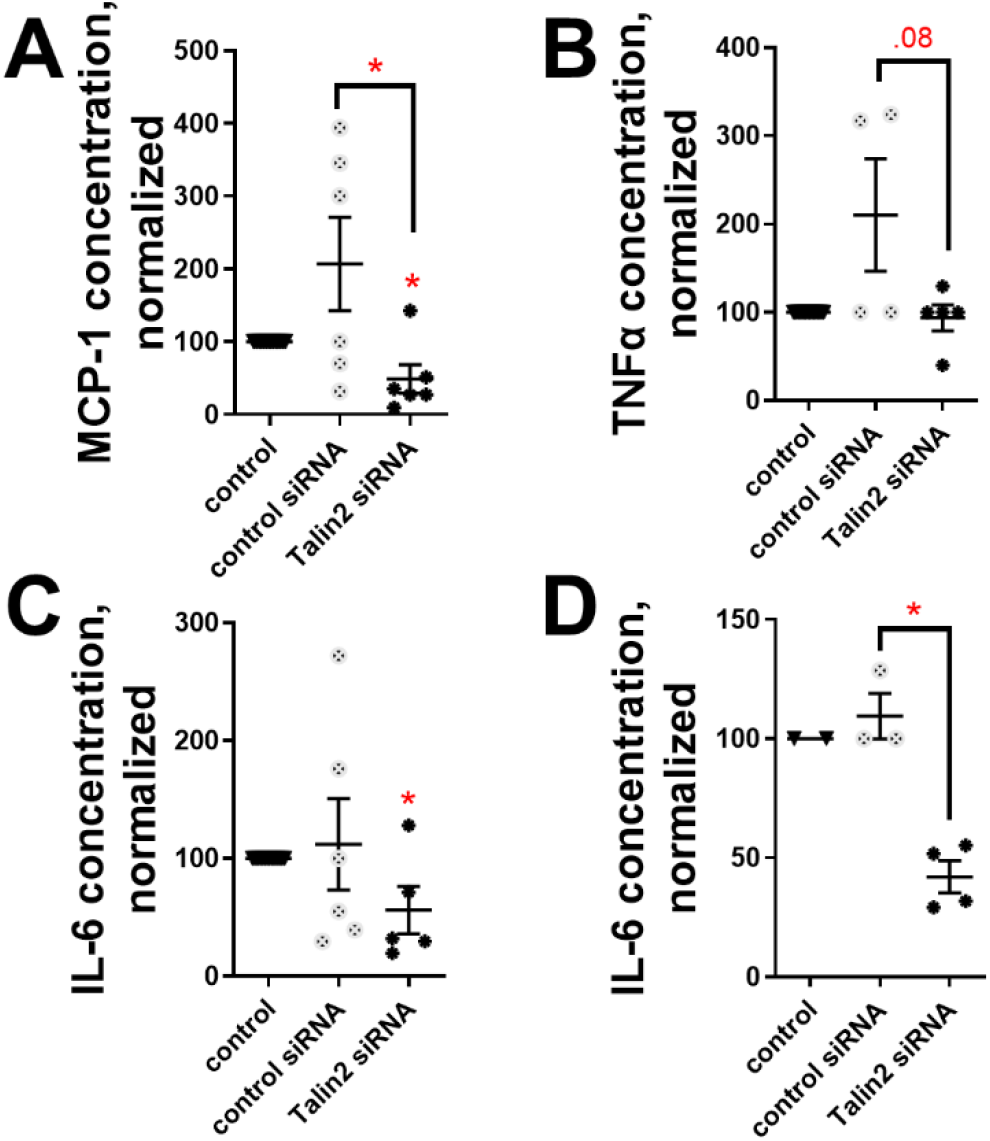
Treatment with talin2 siRNA reduces the pro-fibrotic secretome of human myofibroblasts. Treatment with 50 nM of talin2 siRNA on human monocyte-myofibroblasts significantly lowers the amount of secreted (A) MCP-1 and lowers, though not significantly, the amount of (B) TNF-α. Talin2 siRNA reduced the amount of IL-6 secreted by (C) monocyte-myofibroblasts and (D) fibroblast-myofibroblasts. * = statistical significance of P < 0.05, < 0.01, or < 0.001, significance vs control unless otherwise indicated, Student’s t-test.

**Supplemental Figure 5:**
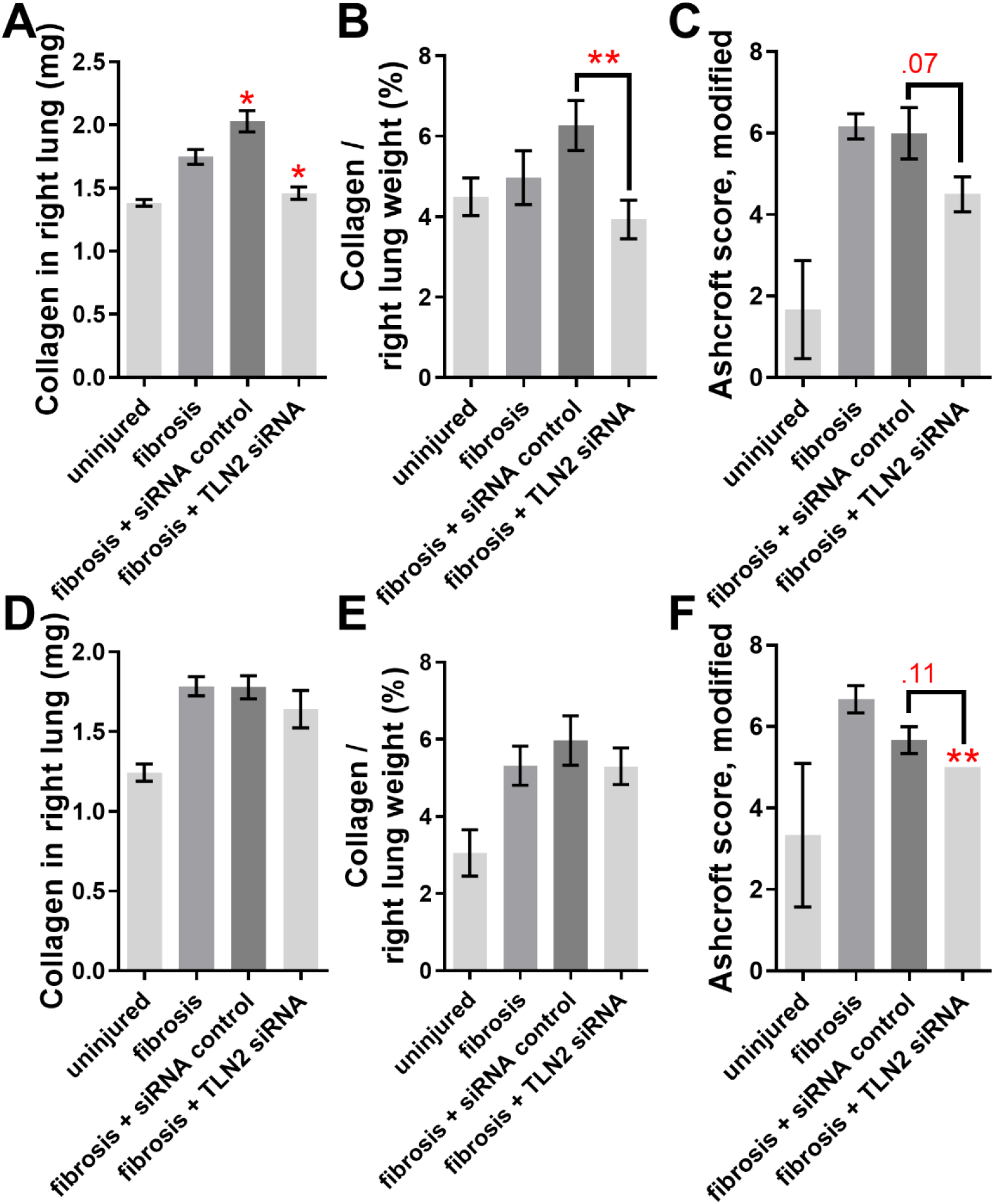
talin2 siRNA rescues the fibrotic damage from both male and female mice. Talin2 siRNA treatment compared across male (n=5 or 6, A, B, C) and female (n=3, D, E, F) mice for amount of collagen in lungs (A, D), collagen as a percentage of dry lung weight (B, E), and blinded Ashcroft scores (C, F). n ranges from 3 to 6. * = statistical significance of P < 0.05, < 0.01, or < 0.001, significance vs fibrotic lungs unless otherwise indicated, Student’s t-test.

**Supplemental figure 6:**
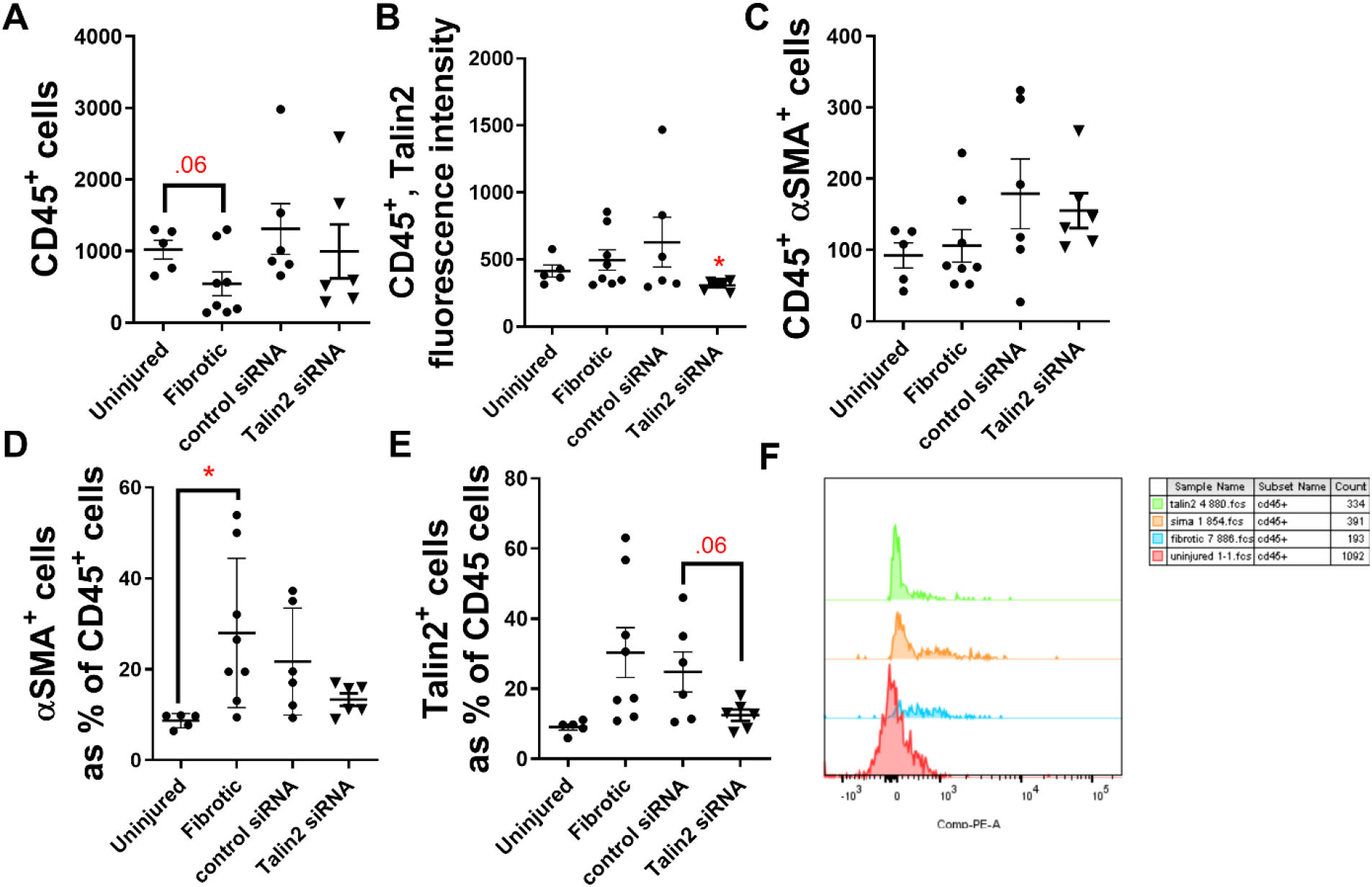
Treatment with talin2 siRNA shows a trend to reduction of the number of CD45^+^ αSMA^+^ and CD45^+^ talin2^+^ cells compared with fibrotic lung in a pilot study. Mouse lungs were insulted with bleomycin, and treated with 0.2 μM of talin2 siRNA. Lungs were lavaged after euthanasia, and assessed for (A) CD45, (B) talin2, and (C) αSMA by flow cytometry. (D) CD45^+^ αSMA^+^ cells normalized to overall CD45^+^ cells from that mouse. (E) CD45^+^ Talin2^+^ cells normalized to overall CD45^+^ cells from that mouse. (F) Representative flow plot of BAL cells. * = statistical significance of P < 0.05, < 0.01, or < 0.001, significance vs control unless otherwise indicated, Student’s t-test.

**Supplemental Figure 7:**
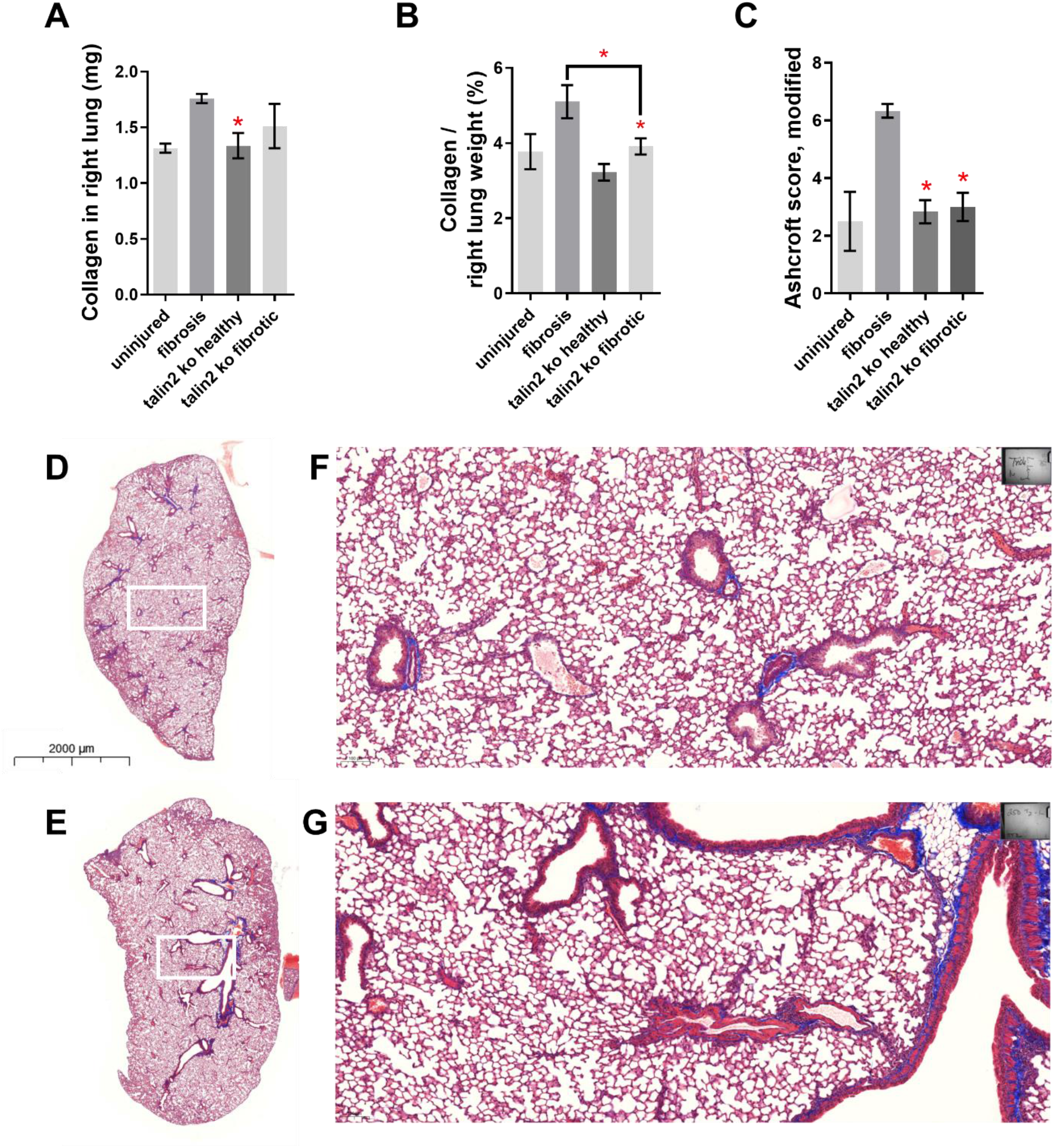
Tln2 -/- mice are resistant to lung fibrosis. Tln2 -/- mice were insulted by bleomycin, and assessed as in Figure 5. (A) Collagen content from the right, multi-lobed lung assessed by hydroxyproline assay. (B) Data from C divided by dry weight of right lobes of mouse lungs. (C) Blinded Ashcroft scoring. (D-E) Representative images of left, single lobed lungs stained with Massons’s trichrome. (F-G) Inset of lungs. (D,F) Uninjured Tln2 -/- lung, (E,G) bleomycin-insulted Tln2 -/- lung. n = 6. * = statistical significance of P < 0.05, < 0.01, or < 0.001, significance vs fibrotic lungs unless otherwise indicated, Student’s t-test.

**Supplemental Figure 8:**
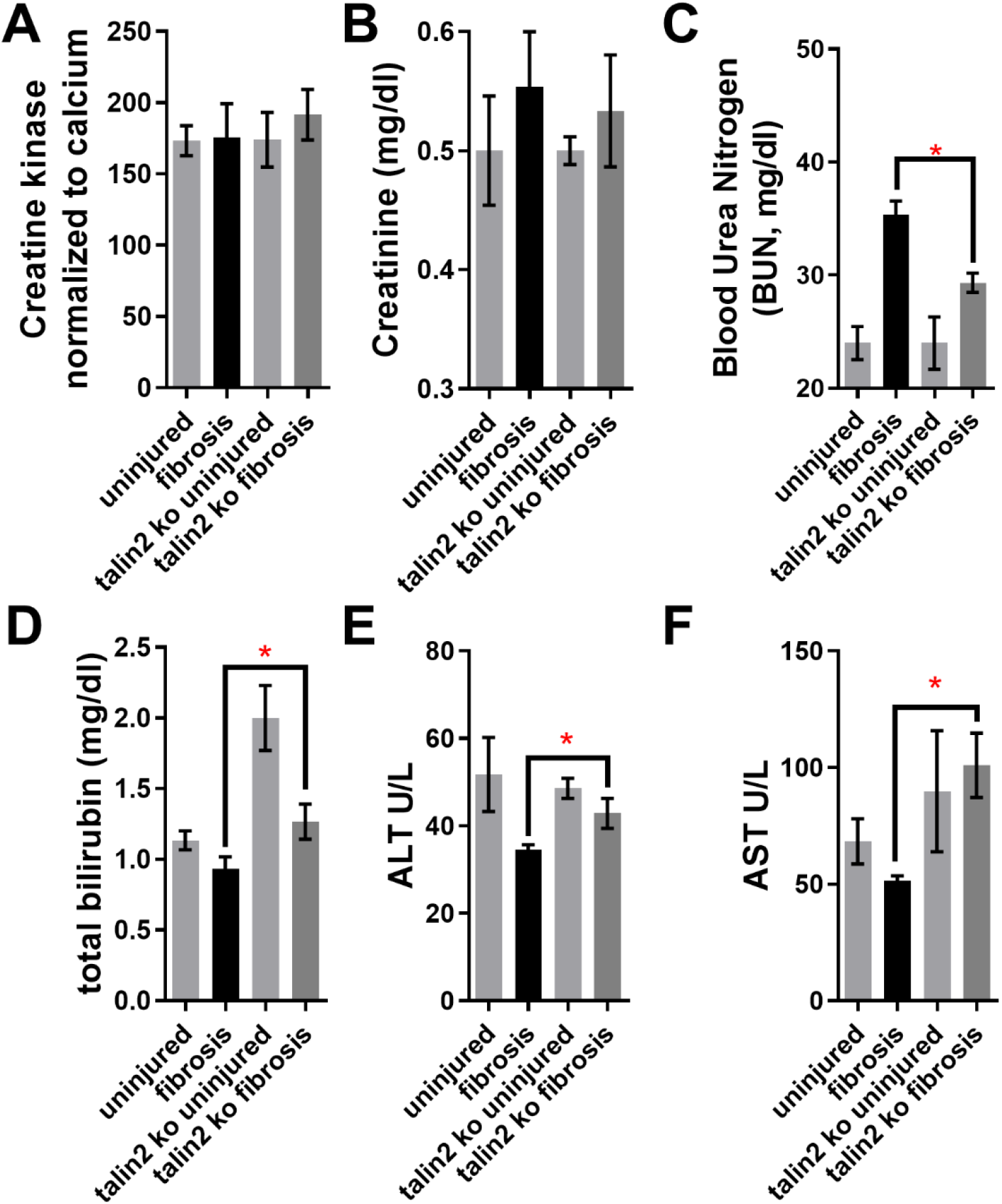
Blood analysis from kidney UUO model. Blood was taken from mice (C57Bl6 and Tln2-/-) directly before sacrifice at 2 weeks post UUO. Concentrations in serum of (A) creatine kinase, (B) creatinine, (C) blood urea nitrogen, (D) bilirubin, (E) ALT, and (F) AST. * = statistical significance of P < 0.05, < 0.01, or < 0.001, Student’s t-test.

**Supplemental Figure 9:**
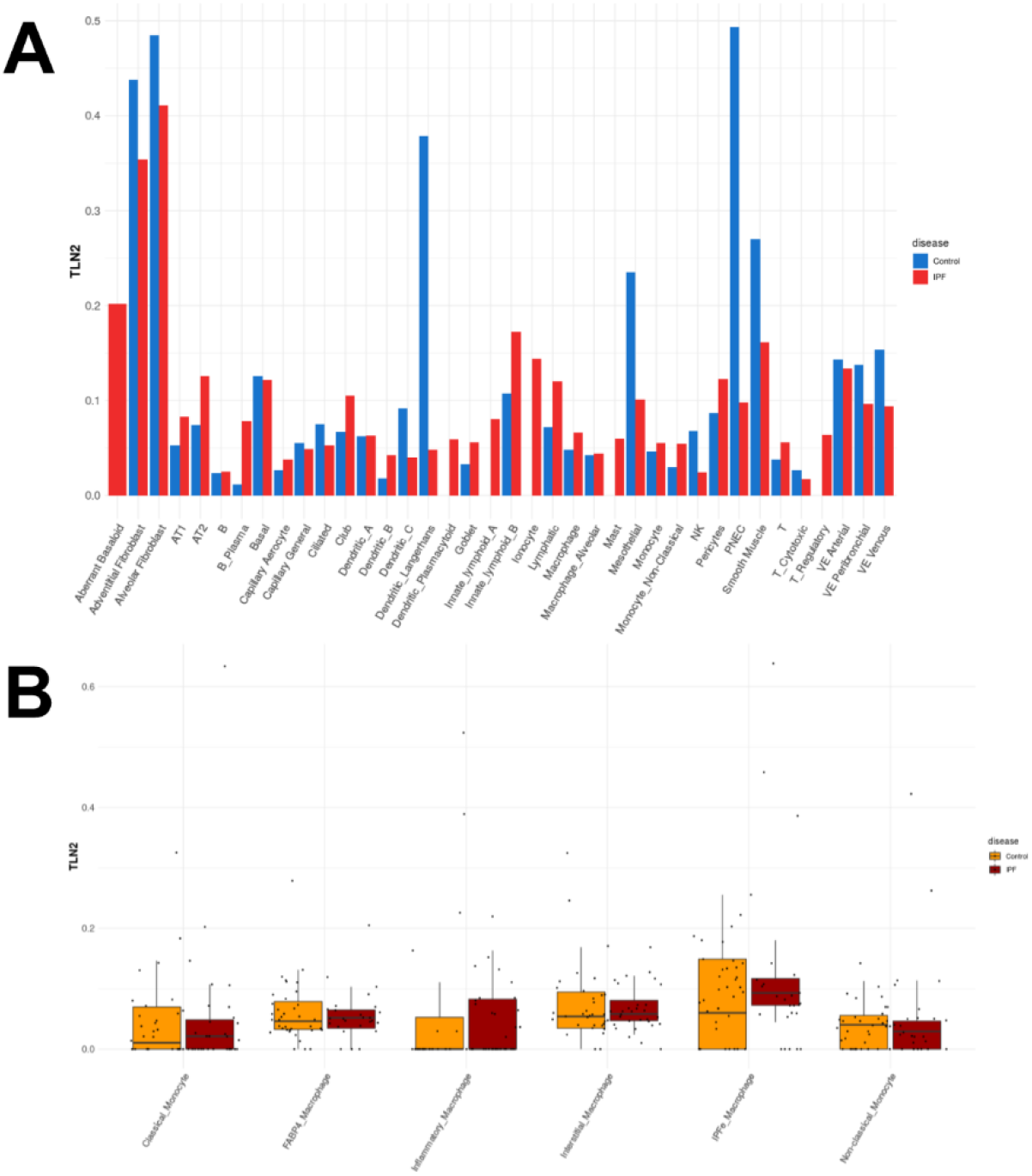
Talin2 transcripts are upregulated among monocyte-derived cell types found in IPF. (A) Talin2 transcript frequency in cells found in IPF. (B) Talin2 transcript frequency by individual patient for selected immune cell subtypes. All data and figures from the IPF Atlas, Kraminski/Rosas dataset.

## Notes

### Competing Interest Statement

MJVW and JH are authors on a patent regarding this work

### Summary of Updates

This version of the manuscript has been updated to include the data from the talin2 -/- mouse, kidney fibrosis data, and to clean up some of the writing for clarity This latest update is the final version of the paper, with some additional knockout data, some minor editing, and removing Roy Zent and David Critchley from the authors. They developed the knockout mouse, but felt their contributions didn't merit authorship, since all they did was share the mouse with me.

